# Learning single-cell spatial context through integrated spatial multiomics with CORAL

**DOI:** 10.1101/2025.02.01.636038

**Authors:** Siyu He, Matthew Bieniosek, Dongyuan Song, Jingtian Zhou, Benjamin Chidester, Zhenqin Wu, Joseph Boen, Padmanee Sharma, Alexandro E. Trevino, James Zou

## Abstract

Cellular organization is central to tissue function and homeostasis, influencing development, disease progression, and therapeutic outcomes. The emergence of spatial omics technologies, including spatial transcriptomics and proteomics, has enabled the integration of molecular and histological features within tissues. Analyzing these multimodal data presents unique challenges, including variable resolutions, imperfect tissue alignment, and limited or variable spatial coverage. To address these issues, we introduce CORAL, a probabilistic deep generative model that leverages graph attention mechanisms to learn expressive, integrated representations of multimodal spatial omics data. CORAL deconvolves low-resolution spatial data into high-resolution single-cell profiles and detects functional spatial domains. It also characterizes cell-cell interactions and elucidates disease-relevant spatial features. Validated on synthetic data and experimental datasets, including Stero-CITE-seq data from mouse thymus, and paired CODEX and Visium data from hepatocellular carcinoma, CORAL demonstrates robustness and versatility. In hepatocellular carcinoma, CORAL uncovered key immune cell subsets that drive the failure of response to immunotherapy, highlighting its potential to advance spatial single-cell analyses and accelerate translational research.

## Introduction

Understanding the spatial organization of cells within tissues is fundamental for deciphering the mechanisms underlying tissue function, development, and diseases. Emerging spatially resolved omics technologies have transformed our ability to study complex tissues by *in situ* measurements of molecular profiles, including gene and protein expression, epigenetic marks, and chromatin structure [1–4]. These innovations have demonstrated tremendous potential in advancing research across diverse fields, including neuroscience [5], cardiovascular pathology [6, 7], cancer biology [2, 8, 9] and developmental biology [10]. The integration of multiple spatial omics modalities offers an exciting opportunity to comprehensively dissect tissue complexity. By combining information across molecular dimensions—such as genes, proteins, and metabolites—spatial omics integration provides a more detailed and nuanced understanding of the intricate microenvironments within tissues.

However, integrating data from distinct spatial omics modalities remains challenging [1]. First, multiomics measurements are often performed on adjacent rather than identical slides, leading to potential discrepancies between cells profiled in different modalities. Second, the spatial resolutions of multiomic measurements are not always consistent. Third, the number of genes, proteins or other molecules measured across different samples can differ. Additionally, the quality of spatial omics data can vary significantly between modalities, further complicating integration efforts. For example, a recent study of glioblastoma [11] used Visium platforms for whole transcriptomic profiling but with spatial spots resolution (each may contain 1-10 cells), along with CODEX platforms for spatial proteomics, which typically measure only tens of proteins at single-cell resolution. This makes the joint single-cell ontology analyses challenging. These challenges underscore the need for methods that can effectively learn a latent representation of the raw measurements that incorporates variability of features from multiple assays, and improve the accuracy and resolution of each modality to provide a more comprehensive and precise biological landscape of complex tissues.

Existing approaches for integrating single-cell multiomics, such as Seurat v4 [12], MOFA+ [13], and totalVI [14], have made significant strides in leveraging multimodal data to provide a unified representation of cellular states and multimodal relationships. However, these methods missed the modeling of spatial contexts, which are critical for understanding cell-cell interactions within their native tissue environments. While advancements have been made in integrating spatially resolved data, many methods focus only on combining histology and spatial transcriptomics [15–17], or only designed for deconvoluting spatial low resolution data referenced on single-cell data such as SpatialScope [18]. SLAT [19] and MIIT [20] primarily address workflows for registering neighboring tissue sections, such as spatial transcriptomics and mass spectrometry imaging, but not leveraging these data to uncover biologically functional tissue regions or infer cellular behaviors. Additionally, models such as MEFISTO [21], SpatialGlue [22] and SpaMosaic [23], provide frameworks for spatial integration but are primarily focused on domain detection, lacking biologically interpretable capabilities and resolution flexibility.

To overcome these limitations, we introduce CORAL (Comprehensive spatial Omics Registration and Analysis for Learning spatial features), a probabilistic, graph-based method designed to integrate diverse spatial omics datasets. Taking multimodality molecular profiles of unmatched spatial resolution and detected features, CORAL generates single-cell embedding with information from both data modalities, deconvolves the lower-resolution modality to infer its profile in individual cells, and predicts interactions between neighboring cells. We validated CORAL using multiple synthetic datasets, spatial CITE-seq datasets from mouse thymus, and paired CODEX and Visium data from hepatocellular carcinoma (HCC) samples treated with anti-PD1 therapy. CORAL successfully delineated the tissue ecosystems in HCC and identified macrophages as key players interacting with CD4+ T cells, mediating immunosuppressive interactions that hinder PD1 immunotherapy responses. Overall, CORAL provides a nuanced and accurate representation of cellular heterogeneity, spatial organization, and underlying molecular mechanisms within complex tissues, offering a powerful framework for the in-depth exploration of spatially resolved molecular landscapes.

## Results

### Overview of CORAL model

CORAL is designed to delineate single-cell spatial contexts by integrating spatial multi-omics data that vary in spatial resolution and molecular features depth (Fig. 1a). The primary objective of CORAL is to predict single-cell resolution molecular profiles within a spatial context by combining the strengths of various spatial omics technologies at cellular resolution or feature throughput. CORAL employs a multimodal approach that captures interactions between molecular layers through a cross-attention mechanism and derives a joint latent representation of cells. Using graph attention layers, the model learns the relationships between neighboring cells, which allows incorporation of spatial dependencies into the latent representations and prediction of cell-cell interactions. Finally, CORAL’s probabilistic framework allows it to integrate data with different spatial resolutions. As a result, it enables the deconvolution of lower resolution spatial data (e.g. Visium) by inferring single-cell expressions from aggregated spot-level data. This is complemented by a flexible variational inference mechanism that models both shared and modality-specific latent variables, thereby preserving the unique information contained within each omics modality while integrating them into a cohesive spatial context (Fig. 1b, Methods). Furthermore, CORAL infers a hierarchical variable associated with spatially variable features (Fig. 1a).

**Fig. 1:**
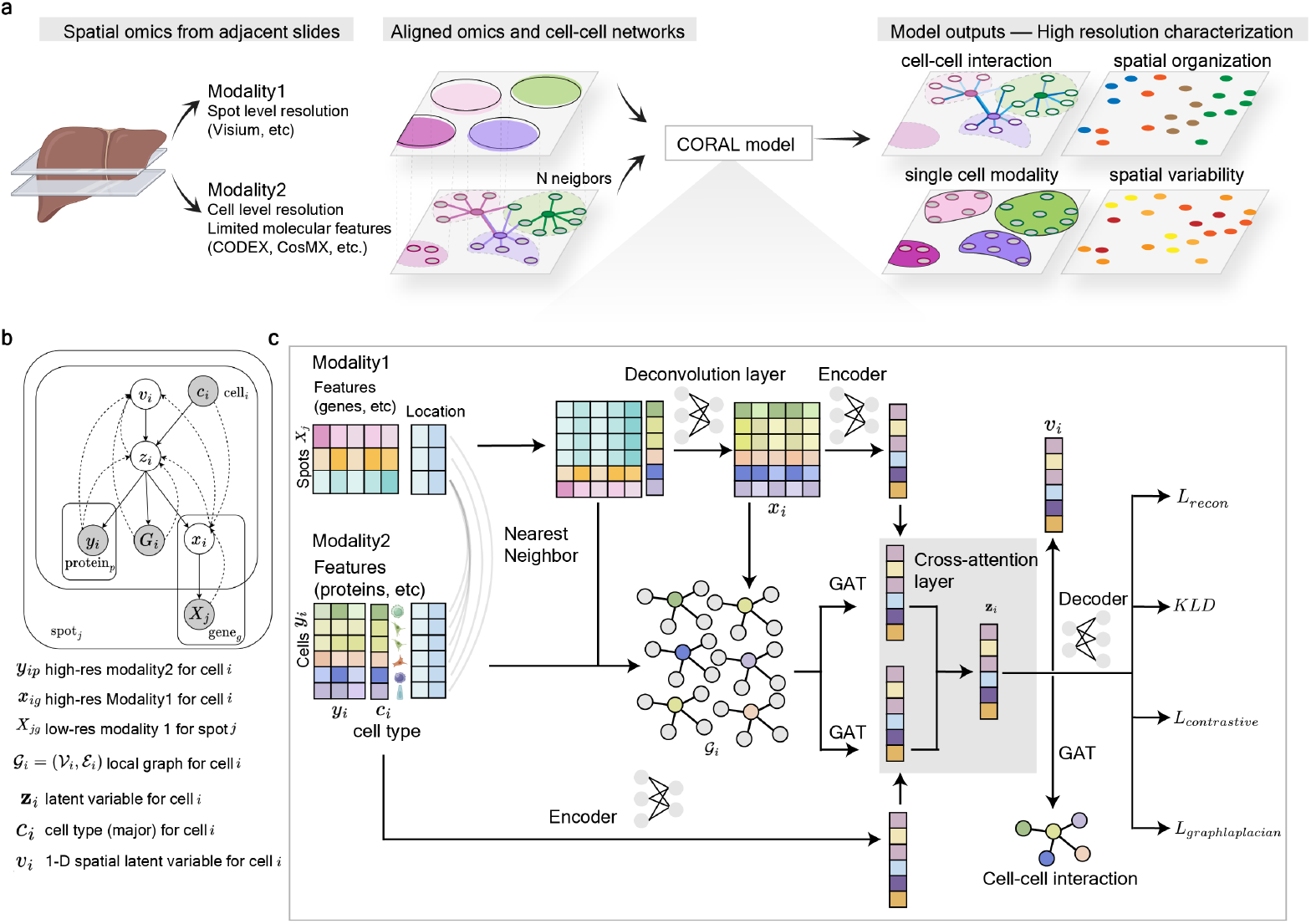
CORAL enables multimodal integration for high-resolution molecular mapping at the single-cell level. **a**, Conceptual overview of CORAL. CORAL integrates spatial omics data across multiple modalities with distinct spatial resolutions. Modality 1 represents lower-resolution spot-based platforms (e.g., Visium), while Modality 2 captures higher-resolution cell-level features (e.g., CODEX). For each individual cell in Modality 2, CORAL aligns spatially adjacent spots from Modality 1. The model reconstructs shared latent variables reflecting the spatial organization of the tissue and performs deconvolution to resolve lower-resolution spot data into cell-level molecular profiles. CORAL further identifies spatially variable features through hierarchies and leverages graph attention networks to model cell-cell interactions within their local environment. **b**, Graphical model of CORAL. Dotted lines indicate the inference process, and solid lines represent the generative process. **c**, CORAL integrates spatial multi-omics data by aligning high-resolution cellular data (Modality 2, e.g., protein expression from CODEX) with neighboring lower-resolution spot data (Modality 1, e.g., gene expression from Visium). The framework first constructs local subgraphs for each cell, incorporating molecular features from its spatial neighbors to capture the local cellular context. The encoder processes these combined features to deconvolute spot-level data, enabling single-cell resolution for gene expression. A graph attention mechanism using graph attention networks (GAT) then models cell-cell interactions, while contrastive learning optimizes the representation of cell types, encouraging similarity among cells of the same type while distinguishing between different types. CORAL outputs include predictions for single-cell transcriptomics, spatial organization, and inferred cell-cell communication, providing a comprehensive molecular map at the single-cell level.

Two aligned modalities are required as inputs, not necessarily sharing the same spatial resolution. When multi-omics data are obtained from different tissue sections, preprocessing steps are typically required. These include cropping overlapping areas, registering tissues, or aligning images using methods like scale-invariant feature transform (SIFT), SLAT [19] or MIIT [20]. The processed data consists of combined expression matrices derived from individual cell locations or spatial spots (Fig. 1c, Methods). Spatial spots represent collections of cells within localized tissue regions due to resolution limitation of the technology, but usually have a wider range of molecules detected. In contrast, cell locations provide higher spatial resolution but are typically limited to a smaller number of molecular features. CORAL also takes coarse annotations of major cell types for single cells (e.g., from CODEX). These annotations could be based on pre-existing labels or simple clustering approaches, such as unsupervised manifold separation. However, due to the limited number of markers available, it often fails to accurately distinguish subtypes or even major cell types. By integrating data from low-resolution modalities with deeper molecular profiling, CORAL refines and enriches cell type annotations, enhancing resolution and providing additional molecular features at the single-cell level.

CORAL begins by mapping each cell to its nearest lower-resolution spot based on spatial proximity. This alignment spatially integrates the datasets and combines molecular profiles from both modalities for each cell. Local sub-graphs centered around each cell were constructed, incorporating data from spatially neighboring cells to capture the local spatial context and interactions among cells. An encoder network processes the combined expression data of these sub-graphs, deconvoluting mixed spot-level gene expression data into single-cell-level expression. This deconvolution allows the model to separate the contributions of individual cell types within each Visium spot. To learn deconvolving, a contrastive loss function was incorporated that encourages cells of the same type to exhibit similar expression profiles while distinguishing them from cells of different types. CORAL also implements a graph attention layer that learns cell-cell interactions within the spatial graph by computing attention weights based on spatial proximity. This mechanism allows the model to focus on local interactions and effectively capture spatial patterns. Furthermore, it includes a layer dedicated to identifying spatial domains (Methods). This layer captures higher-level spatial structures by aggregating information across cells and their neighbors, enabling the model to recognize and characterize spatial domains - regions within tissue that share common molecular features and neighboring cell identities. By detecting these spatial domains, CORAL enhances the understanding of tissue organization and function, providing insights into how cells interact within their microenvironments and contribute to overall tissue architecture. To optimize its performance, reconstruction losses were utilized to tailored to each modality and enforces spatial smoothness through Laplacian regularization while modeling complex dependencies via Kullback-Leibler divergence terms. This strategy allows CORAL to leverage the strengths of different spatial omics modalities, providing a comprehensive and high-resolution view of the molecular landscape at single-cell resolution.

To evaluate it’s performance in spatial domains detection and spot deconvolution, we first compared it against state-of-the-art methods (totalVI, COVET, SpatialGlue, and spatialScope) using both synthetic and real datasets. These datasets included synthetic data provided by SpatialGlue, truncated and downsampled MERFISH mouse cortex data, and synthetic spatial CITE-seq data produced with scDesign [24]. We then demonstrated CORAL’s utility through two in-depth case studies. In the mouse thymus, it effectively characterized tissue structures and tracked immune cell maturation. In human hepatocellular carcinoma, it delineated immune-tumor interactions, uncovering key ecosystem features that contribute to the suppression of immunotherapy responses.

### Coral enables integrated single-cell spatial context delineation for synthetic multi-omics data

We evaluated CORAL across four key functionalities: predicting spatial domains, deconvoluting spot-resolution modalities into single cells, identifying spatial variables factors, and predicting cell-cell interactions. While no method currently achieves all four functionalities simultaneously, we benchmarked each functionality against similar methods to validate efficiency and accuracy.

To assess the ability to identify spatial domains using integrated multimodal data, we first used the simulated dataset in SpatialGlue that included both transcriptomic and proteomic profiles [22]. To further simulate the spatial resolution differences between modalities, we intentionally reduced the spatial resolution of one modality (modality 1) by aggregating the neighboring cells (Supplementary Fig. 1a). We evaluated CORAL’s performance by comparing it with several baselines: modality 1 alone, modality 2 alone, the downsampled modality 1 alone, concatenated PCA of both modalities, SpatialGlue, and totalVI. For this experiment, we processed the downsampled modality 1 alongside the original modality 2. In contrast, methods like SpatialGlue and TotalVI [14] that require both modalities to be at the same resolution were run with the original high-resolution data. CORAL demonstrated comparable performance to SpatialGlue in detecting spatial patterns and background spatial factors (Supplementary Fig. 1b-c), as validated by multiple performance metrics, including homogeneity, mutual information, V-measure, and adjusted mutual information (AMI). Both SpatialGlue and CORAL, which incorporate spatial neighbor information and attention layers, significantly outperformed models like TotalVI, which relies solely on single-cell resolution data (Supplementary Fig. 1d). Proper integration of two modalities greatly enhances the detection capabilities of spatial domains and cell-cell interactions. In addition to domain detection, CORAL also improves the spatial resolution of downsampled modalities. By comparing the deconvoluted gene expression from the downsampled modality 1 with the original high-resolution modality 1, we observed a high accuracy (Supplementary Fig. 1e-g).

To further explore model’s single-cell analysis capabilities, we applied it to a synthetic MERFISH dataset of the mouse primary motor cortex [25], previously analyzed by SpatialScope [18]. This dataset captures the laminar organization of the cortex, served as a ground truth for spatial organization. We simulated two different modalities: first, by downsampling the MERFISH data (Fig. 2a, Supplementary Fig. 2a) noted as synthetic modality 1, and second, by truncating the gene set from 254 to 50 genes (synthetic modality 2). We compared CORAL’s performance against Spatial-Glue and COVET[26] (Supplementary Fig. 2b-d) on spatial domain detection with multiple downsampling ratio of modality 1, and found that (Fig. 2a-b), CORAL consistently outperformed synthetic modality 2 and SpatialGlue across all fours different magnitude of sampling, and outperformed COVET with the highest magnitude of our sampling (with grids number 5*×*15) (Fig. 2b). Additionally, CORAL successfully imputed single-cell resolution expression data from the downsampled modality, highly correlated with the ground truth (Fig. 2c), and performed better than SpatialScope [18], reinforcing it’s efficacy in reconstructing single-cell expression from low-resolution modalities.

**Fig. 2:**
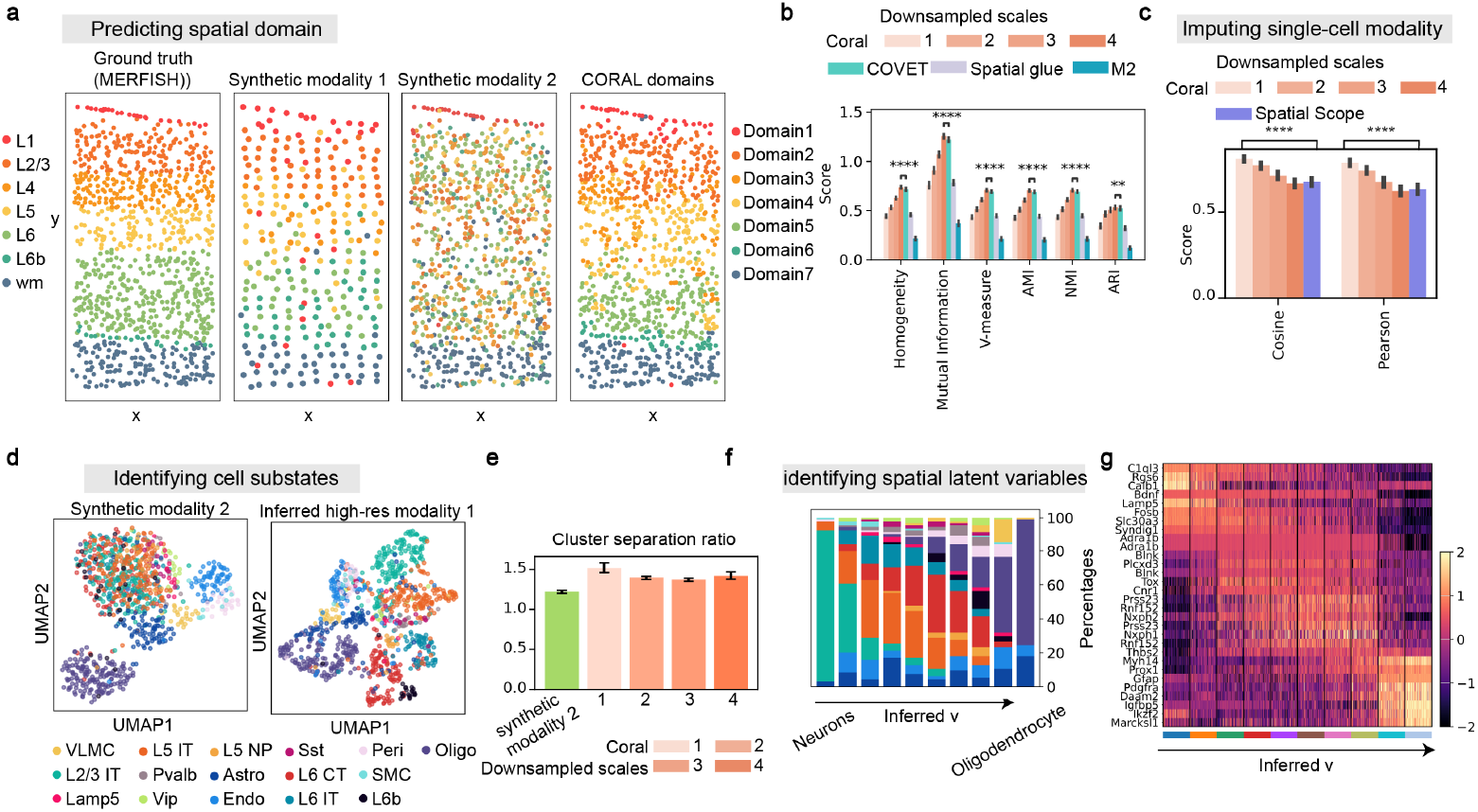
CORAL maps single-cell spatial context using truncated and down-sampled MERFISH data. **a**, Spatial domains identified by CORAL, synthetic modality 1 and synthetic modality 2 (colored by identified domains), compared to ground truth (colored by cortical layers) in the mouse motor cortex. **b**, Performance metrics (Homogeneity, Mutual Information, V-measure, Adjusted Mutual Information (AMI), Normalized Mutual Information (NMI), Adjusted Rand Index (ARI)) comparing CORAL’s spatial domain detection accuracy at varying levels of down-sampling with alternative methods (COVET, Spatial Glue) and truncated MERFISH data (Modality 2). A higher value for these metrics indicate better performance. The downsampled scales (1,2,3,4) correspond to grid sizes of 20*×*60, 15*×*45, 10*×*30, and 5*×*15 respectively, representing an average of 1.5 cells, 2 cells, and 3.5 cells, and 12.5 cells per spot for synthetic modality 1 (Supplementary Fig. 2). CORAL consistently outperformed synthetic modality 2 and SpatialGlue across all fours different magnitude of sampling, and outperformed COVET with the highest magnitude of our sampling (with grids number 5*×*15) Two-sided independent two-sample t-test was performed on bootstrapped scores between SpatialGlue and CORAL (n=10), P values = 5.30 × 10^*−*26^, = 9.55 × 10^*−*23^, = 1.48 × 10^*−*17^, = 1.39 × 10^*−*17^, = 1.48 × 10^*−*17^, = 4.89×10^*−*3^ respectively. **c**, Similarity scores (Cosine similarity and Pearson correlation) between CORAL’s predictions and Spatial Scope across different downsampling scales. Coral outperformed Spatial Scope across scale 1-3 though worse at scale 4 (downsampling is 12.5 cells per spot for synthetic modality 2). A higher value for these metrics indicate better performance. One-way anova test was performed on bootstrapped testing (n=10), P values = 2.54*e −* 76 and = 7.64*e −* 90 separately, ****P*<*0.0001, **P*<*0.01. **d**, UMAP projections of synthetic modality 2 (Left) and inferred high-resolution/single-cell (Right) MERFISH data, with points colored by annotated major cell type. **e**, Cluster separation ratio (Methods) across clusters of synthetic modality 2 and CORAL imputed high-res modality 2. A higher value for this metric indicate better performance. It indicated CORAL successfully distinguished various cell types and subtypes from the deconvoluted modality 1. **f**, Stacked bar plot showing cell type proportions across a gradient of the spatial latent variable *v*, indicating a shift from neuron to oligodendrocyte prevalence. The color of bars is the same as that in panel d. **g**, Heatmap of expression of variables genes along inferred *v*, further indicating the gene signature shifting from Neurons-like to oligodendrocytes.

Furthermore, CORAL excels in identifying cell substates from truncated or low-resolution data. For example, in modality 2 where the number of profiled genes were limited, the capabilities to identify the fine cell states is limited (Fig. 2d). It successfully distinguished various cell types and subtypes from the deconvoluted modality 1, including subtypes of excitatory and inhibitory neurons, as well as other nonneuronal cells, suggesting the importance of taking advantage of multiple data modalities (Fig. 2d-e). The one-dimensional latent variable *v* compresses their molecular profiles and spatial contexts. In the MERFISH data, it corresponding to the change of percentages of neurons (Fig. 2f), along with the gene signature (Fig. 2g). By projecting cellular measurements to *v* latent space, CORAL reveals underlying spatial patterns that may not be apparent from gene space. Finally, to demonstrate that our method can capture spatial networks, we predicted cell-cell interactions in cortical layers(Supplementary Fig. 3). By integrating spatial proximity and molecular data, CORAL quantified interactions between cell types in mouse brain, such as astrocyte-neuron interactions, enabling potential comparisons of the functional dynamics of the tissue across samples [27].

### CORAL integrates RNA and protein data to refine tissue domains and resolve single-cell heterogeneity

Besides directly downsampling RNA features, we also evaluate CORAL’s performance on more realistic paired single-cell RNA and protein data. We used scDesign3[24] to simulated CITE-seq data based on a reference dataset [28] and assigned the different cell types to different spatial regions with defined probabilities (Supplementary Fig. 4). We also downsampled cells for the simulated RNA modality to mimic the different spatial resolutions between modalities (Fig. 3a). The simulated datasets kept the original data structure of CITE-seq (e.g. gene-gene correlation, cell-cell distance), and provide gold standard of cell types and associated spatial domains to benchmark the performance of different models on spatial domain detection and cell-type classification. CORAL outperformed other methods, including SpatialGlue, in classifying different cell types and map their associated spatial domains. For instance, CORAL distinguished the spatial regions assigned to plasmacytoid dendritic cells (pDCs) and dendritic cells (DCs), which cannot be identified with either modality without incorporating spatial information (Fig. 3b-c). Furthermore, CORAL effectively imputed low-resolution RNA data into single-cell resolution (Fig. 3d, Supplementary Fig. 4e). This imputation addresses the challenge of cell-type identification that occurs in protein data with a smaller number of features (Fig. 3e-f). Notably, the imputed single-cell RNA data identified cell types that aligned with the domain detection results, showing consistency in both tasks.

**Fig. 3:**
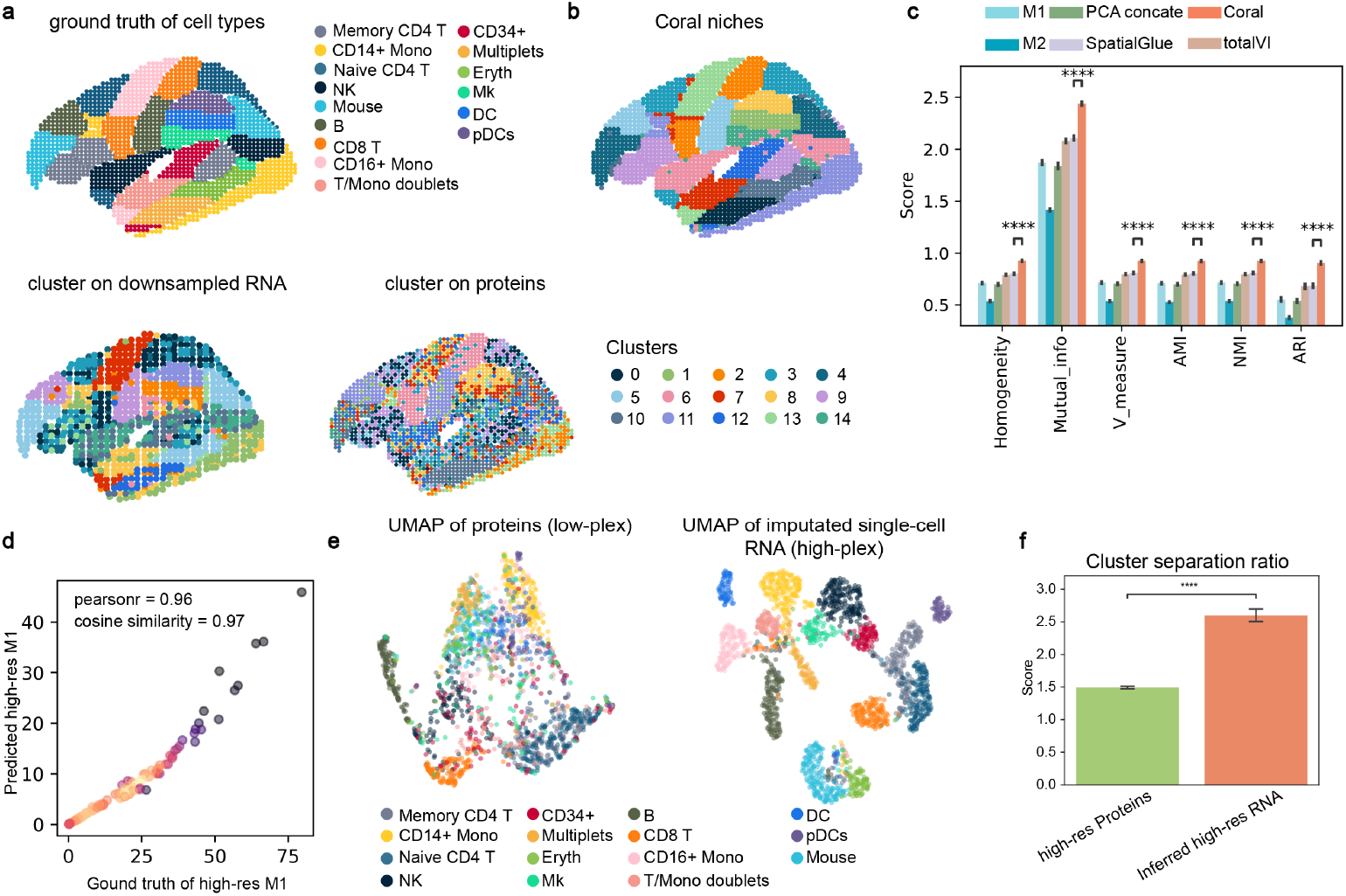
CORAL delineates single-cell spatial context in simulated spatial CITE-seq data. **a**, Upper: Ground truth of cell types sampled from CITE-seq data with immune cells forming domains as brain regions. Bottom: Cluster patterns identified on downsized RNA-sequencing, and proteins. The colors are clusters with the same number as the ground truth of cell types. **b**, Spatial domains identified by CORAL model, the colors are clusters with the same number as the ground truth of cell types. **c**, benchmarking the spatial domains detction across modality 1 (RNA), modality 2(protein), PCA of two modalities, Spatial Glue, totalVI and Coral. All the number of domains are equal to the ground truth of cell types. A higher value for these metrics indicate better performance. CORAL outperformed all other methods, including SpatialGlue, in classifying different cell types and map their associated spatial domains. Two-sided independent two-sample t-test was performed on bootstrapped scores between SpatialGlue and CORAL (n=10), P values = 1.47 × 10^*−*17^, = 1.66 × 10^*−*18^, = 4.25×10^*−*17^, = 4.18×10^*−*17^, = 4.26×10^*−*17^, = 4.02×10^*−*17^ respectively.****P*<*0.0001. **d**, scatter plot of ground truth of single-cell modality 1 (RNA) and deconvoluted modality 1, the color represented the density of the scatter plot. **e**, UMAP of modality (proteins, low-plex) only and the imputated single-cell RNA (high-plex), colored by ground-truth cell types. **f**, Intra-to-inter-cluster distance ratio between high-res proteins and inferred high-res gene expression. Two-sided independent two-sample t-test was performed. A higher value for this metric indicate better performance. The imputation of CORAL addresses the challenge of cell-type identification that occurs in protein data with a smaller number of features. P value = 4.0 × 10^*−*18^. ****P*<*0.0001.

### Dissecting fine structures and immune cell maturation landscape in mouse thymus through CORAL

To demonstrate CORAL’s broad applicability to real data across a wide spectrum of technology platforms, we applied it to Stereo-CITE-seq data from mouse thymus section [29]. The thymus plays a critical role in the development and maturation of T cells (thymocytes), and its architecture consists of lobules enclosed by a capsule, with connective tissues separating the lobules (Fig. 4b)[30]. Stereo-CITE-seq enables the measurement of both mRNA and protein at subcellular resolution, which provides a gold standard to evaluate CORAL’s capability to impute high resolution data (Fig. 4a). To note, it’s not practical to measure full resolution always, thus a method for resolution augmentation is necessary. We downsampled the RNA data to a lower resolution and test if the joint analysis with CORAL could recover the high-resolution profiles. Clustering with only one modality cannot capture the fine structures of the thymus - protein data resulted in less accurate clustering and the downsampled RNA data produced clusters at a lower resolution (Fig. 4a). In contrast, CORAL improved the resolution of the RNA modality, refining regions that could only be identified with the protein modality, such as the medulla, connective tissue structures, and the multilayered cortex. CORAL’s domain identification capabilities were comparable to those of SpatialGlue (Fig. 4c) [22] despite working with lower resolutio input data, CORAL performed nearly as well as full-resolution SpatialGlue. Additionally, CORAL achieved the highest Moran’s I score, indicating higher correlation between cell embeddings in spatial proximity (Fig. 4d).

**Fig. 4:**
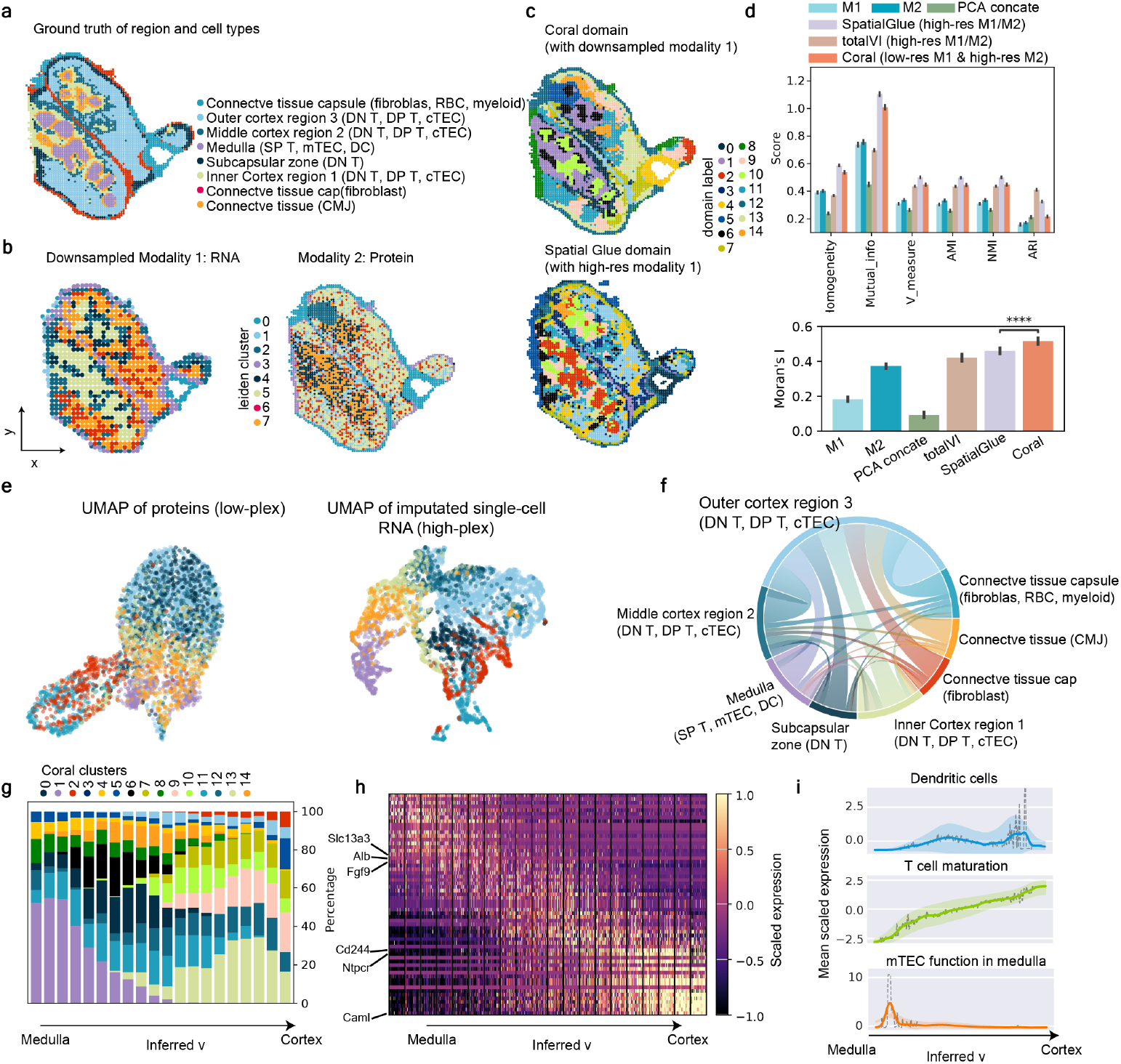
Coral dissects fine structures and gradient of immune cell maturation in mouse thymus. **a**, Expert-annotated region and cell types in mouse thymus. **b**, Leiden cluster of downsampled modality 1: gene expression and modality 2: protein expression in Stereo-CITE-seq data. **c**, Leiden clustering of spatial latent variables identified by CORAL and SpatialGlue. The resolution of clustering was adjusted to make same number (n=15) of clusters on CORAL and SpatialGlue samples.s. **d**, Upper: benchmarking the spatial domains detection with metrics (Homogeneity, Mutual Information, V-measure, Adjusted Mutual Information (AMI), Normalized Mutual Information (NMI), Adjusted Rand Index (ARI)) across modality 1 (RNA), modality 2(protein), PCA of two modalities, Spatial Glue, totalVI and Coral. All the number of domains are equal to the ground truth of cell types. A higher value for these metrics indicate better performance. Bottom: Moran’s I scores for each method. Two-sided independent two-sample t-test was performed between SpatialGlue and CORAL. P value=1.07×10^*−*6^. A higher value for Moran’s I scores indicates better performance. **e**, UMAP of modality (proteins) only and the deconvoluted RNA, colored by ground-truth cell types. **f**, Circular plot of cell-type interactions, where bar lengths represent total mean interactions and band widths represent interaction strength of cell-cell interactions across ground-truth regions of mouse thymus. CORAL suggested strong interactions between cells in the outer cortex region and other regions of the thymus. **g**, Stacked bar plot showing cell type proportions across a gradient of inferred *v*, indicating a shift from medulla to cortex prevalence. The color of bars indicates the CORAL cluster. **h**, Heatmap of scaled expression of variables genes along inferred *v*, indicating gene signatures shifting from medulla to cortex. **i**, Lineplot of gene signatures related to dentric cells, T cell maturation, and mTEC functiosn in medulla along with the inferred *v*.

The comparison of CORAL’s deconvoluted single-cell mRNA expression with ground truth further confirmed its ability to improve the capabilities and resolution of the modalities (Fig. 4e). The predicted single-cell RNA data exhibited a high accuracy when compared to the ground truth gene expression (Supplementary Fig. 5). Furthermore, CORAL suggested strong interactions between cells in the outer cortex region and other regions of the thymus(Fig. 4f). The one-dimensional spatial latent variable *v* captures the transition from the medulla to the cortex in the mouse thymus (Fig. 4g). We identified genes whose expression correlates with *v*, and found that many play key roles in thymic development and T cell maturation. For exampe, the metabolite transporter gene *Slc13a3* is involved in regulaing cellular metabolism within the thymus [31]. Additionally, the *Alb* gene, expressed by medullary thymic epithelial cells (mTECs), functions as a tissue-restricted antigen in the thymus. This expression contributes to the negative selection of self-reative T cells, which is essential for maintaining immune tolerance and preventing autoimmunity [32, 33]. The gene *Fgf9* plays a role in the maturation and structural development of thymic epithelial cells, supporting thymic architecture and function [34, 35]. Furthermore, *Cd244* and *Caml* are both linked to the maturation of thymocytes. *Cd244* is involved in immune cell signaling, while *Caml* regulates calcium signaling, a critical process in thymocyte development and function [36–38] (Fig. 4h). Consistently, the increased expression of gene signatures related to T cell maturation has been identified along with the gradient of *v*, as well as the fluctuation of dendritic cell, and mTEC function in medulla decreased, indicated the interpretability of *v* (Fig. 4i).

### Uncovering tumor microenvironment in hepatocellular carcinoma integrating CODEX and Visium with CORAL

To show how CORAL can recover clincally relevant biological signatures, we applied CORAL to integrate and analyze CODEX and Visium data from two hepatocellular carcinoma patients undergoing immune checkpoint therapy (Supplementary Fig. 6). The dataset includes pretreatment biopsies and post-treatment samples from tumors of two patients: P73 (a non-responder) and P84 (a responder). The CODEX platform features immunofluorescent imaging of 51 markers at subcellular resolution, enabling the identification of major cell types such as tumor cells, CD4+ T cells, macrophages, and B cells. In contrast, Visium provides broad poly-A capture at spot-level resolution, with each spot representing multiple cells. Each of these separate assays has technical limitations that make it difficult to comprehensively understand important tissue microenvironments. By integrating these modalities, CORAL enhanced the resolution of spatial data, enabling us to dissect the tumor microenvironment and identify regulatory mechanisms underlying the differential responses to immunotherapy. Using CORAL, we first identified data-driven spatial domains reflecting tissue organization in both modalities (Fig. 5a). These domains encompassed major cell types across the samples (Supplementary Fig. 7a-b) and were further classified into three categories: intratumoral domains, immune domains, and stromal domains. Intratumor domains exhibited sample-specific characteristics. For example, domains labeled tumor-1 and tumor-2 corresponded to pretreatment P73, while tumor-4 was associated with posttreatment P73. Similarly, tumor-3 and tumor-7 were unique to pretreatment P84, while tumor-6 was specific to post-treatment P84. Tumor-5, however, was shared between P73 and P84, suggesting common spatial and molecular features in these HCC samples.

**Fig. 5:**
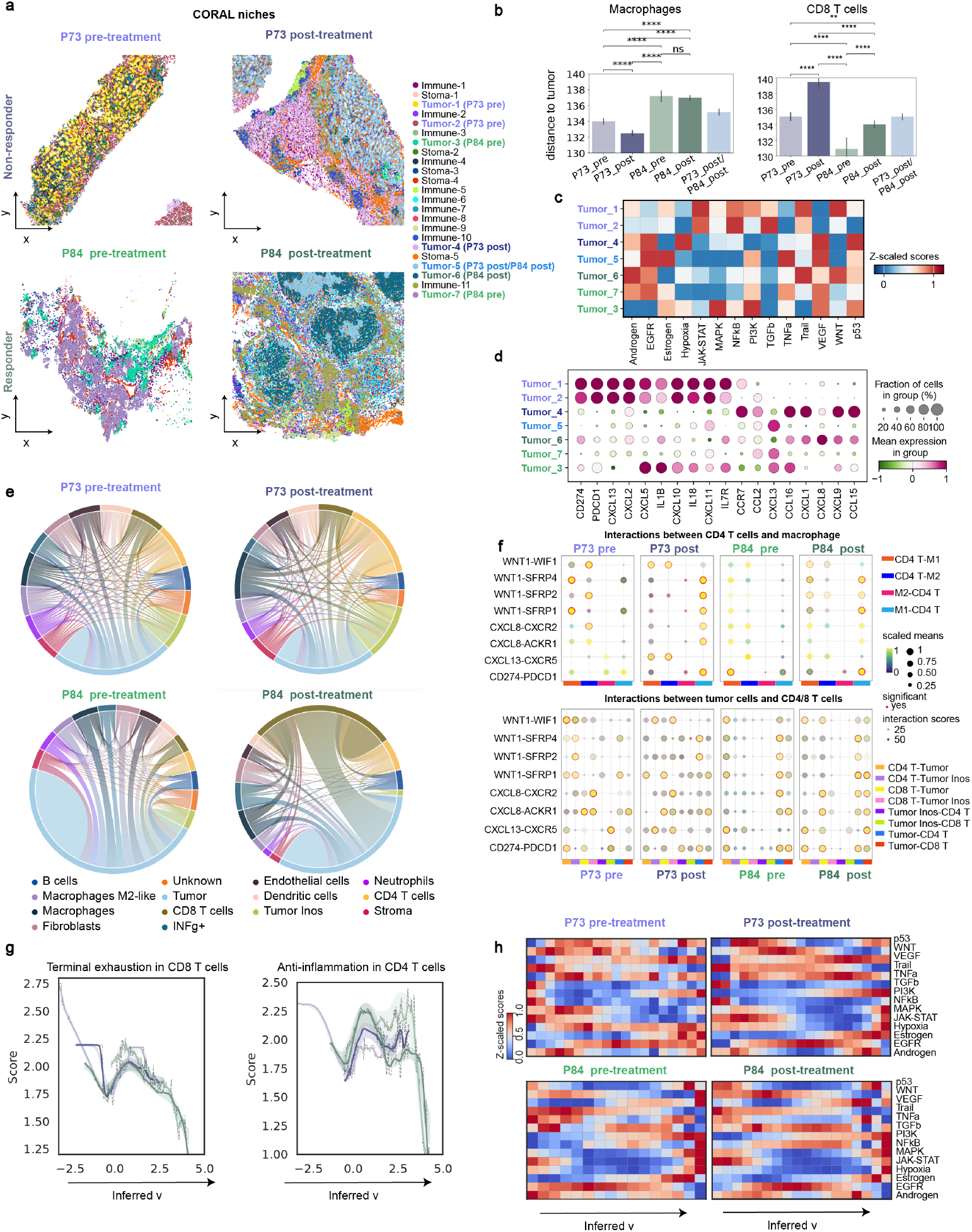
Uncovering the tumor microenvironment in hepatocellular carcinoma by integrating CODEX and Visium through CORAL. **a**, Spatial domains in hepatocelluar carcinoma identified by CORAL using CODEX and Visium data across four samples (P73 pre-treatment, P73 post-treatment, P84 pre-treatment, P84 post-treatment). CORAL identified 11 immune domains, 7 intratumor domains, and 5 stromal domains across these samples. **b**, Distance of macrophages and CD8+ T cells tumor cells within intratumor domains. Intratumor domains are classified as P73-pre, P73-post, P84-pre, P84-post, and shared domains (P73-post/P84-post). A two-sided independent two-sample t-test w1a3s performed. **c**, Z-scaled scores for signaling pathways across intratumor domains. **d**, Z-scaled expression of cytokine genes across intratumor domains. **e**, Interaction score between major cell types across samples. **f**, Scores of ligand-receptor interactions between CD4+ T cells and macrophages across samples, computed using cellphoneDB. **g**, Scores of terminal exhaustion in CD8+ T cells and anti-inflammatory activity in CD4+ T cells, along with inferred parameter *v*. **h**, Z-scared scored of signaling pathways, along with inferred *v* across samples.

Immune domains quantified distinct immune cell distributions. For example, immune-10, predominantly composed of CD4+ T cells and noticed as lymphoid aggregates, was uniquely present in post-treatment P73 as lymphoid aggregates, while immune-2, enriched with macrophages, was scattered across all samples (Fig. 5a). Quantitative analysis of immune cell distances captured that macrophages were significantly closer to tumor cells in the non-responder sample (P73) compared to the responder sample (P84) (Fig. 5b). Conversely, CD8+ T cells were spatially excluded in P73, indicating an immune-evasive tumor microenvironment. Notably, macrophageinfiltrated domains, such as tumor-4 in P73, exhibited recruitment of CD4+ T cells, further contributing to an immunosuppressive milieu. Cytokines are the major immune recruitment factors, and typically are difficult to measure using in situ stains. Visium transcriptomics can quantify these factors, but not identify their cell of origin. CORAL also imputed single-cell-resolved transcriptomic data, effectively increasing resolution of downstream analysis (Supplementary Fig. 7c-d). Thus it facilitated a detailed investigation of cytokine expression(Fig. 5c), revealing heterogeneity across intratumor regions. In pretreatment P73, pathways such as JAK-STAT signaling, which promotes the aberrant synthesis of PD-L1 and facilitates immune resistance, were significantly enriched, linking this pathway to poor responses to PD-1 blockade. Additionally, TGF*−β*, a negative prognostic signal and predictor of immune checkpoint inhibitor resistance, was upregulated in P73 [39, 40]. WNT signaling pathways, also enriched in P73, were implicated in macrophage recruitment [41]. CORAL further facilitated the investigation of cytokine expression, which is typically not fully measurable with CODEX (Fig. 5d). Notably, pretreatment P73 (non-responder) exhibited enrichment of CD274 (PD-L1), PDCD1 (PD-1), and chemokines such as CXCL13 and CXCL12, suggesting a tumor-driven mechanism for macrophage migration and immune evasion[42]. The intratumor regions for post-treatment non-responder, showed enrichment of CCR7, CCR1, CXCL9 and CCL15. In contrast, the post-treatment of responder (“Tumor 6”) and a shared region “Tumor 5” showed enrichment of CXCL8 and preteatment was associated with CXCL3. Notably, CXCL8 is a promising potential prognostic marker for hepatocellular carcinoma [43].

To investigate how immune cells near intratumoral regions shape the tumor microenvironment in non-responder and responder samples, we examined the interactions between key cell types provided by CORAL Fig. 5e). Interestingly, cell-cell interaction predictions by CORAL indicated high interaction between CD8+ T cells and tumor cells in responder (P84), which is indicative of an effective immune response. This kind of interaction is commonly associated with successful immunotherapy, where CD8+ T cells kill tumor cells[44]. In contrast, P73 (non-responder) had more interactions between CD4+ T cells and other immune cells. CD4+ T cells can be regulatory (Tregs) or inflammatory (Th1, Th2), but when they are more involved in interactions in tumors, it often means there is immune suppression[45]. Additionally, macrophages in P73 also interacted more with CD4+ T cells, which further points to an immunosuppressive environment that supports tumor survival[46, 47]. In summary, the non-responder (P73) shows a more immunosuppressive tumor microenvironment, with CD4+ T cells and macrophages that suppress anti-tumor immune responses, while the responder (P84) has stronger CD8+ T cell-mediated immune activity.

The improved resolution of Visium by CORAL allowed for the investigation of specific ligand-receptor interactions that underpin immune cell communication within the tumor microenvironment using CellPhoneDB[48] Fig. 5f, Supplementary Fig. 8). Interestingly, in the non-responder (P73), WNT1/SFRP2 and CXCL13/CXCR5 interactions were significantly enriched between CD4+ T cells and macrophages. Among the interactions between tumor cells and CD8+ T cells, we found CD274/PDCD1 (PD-L1/PD-1) interactions to be significantly enriched in both pre- and post-treatment stages in P73. This suggests the PD1 signaling persists even after blockade that hinder CD8+ T cell-mediated cytotoxicity, a hallmark of resistance to immunotherapy[49]. In contrast, P84 (responder) exhibited lower PD-L1/PD-1 interactions post-treatment, highlighting a reduction in immune evasion and a more favorable immune environment post-therapy. Strong WNT1/SFRP1 and WNT1/SFRP4 interactions were observed in P73, potentially involved in WNT signaling, which has been implicated in recruiting immunosuppressive macrophages and contributing to CD8+ T cell exclusion[50]. This is consistent with findings that WNT signaling can enhance the macrophage polarization toward an M2 phenotype, which is immunosuppressive and promotes tumor progression[51]. Additionally, CXCL8/CXCR2 and CXCL8/ACKR1 interactions enriched in P73 as pro-inflammatory chemokine and tumor progression pathways [52]. CORAL’s inferred latent variable (*v*) captured spatial heterogeneity and functional activity within individual samples (Fig. 5g-h,Supplementary Fig. 9). In P73, CD8+ T cells exhibited exhaustion, characterized by sustained inhibitory signals post-treatment, suggesting a dysfunctional phenotype. Additionally, an enriched antiinflammatory signature in CD4+ T cells at low-*v* regions in P73 pointed to immune dysregulation, further contributing to ineffective responses to therapy.

Overall, CORAL enabled the exploration of key immune escape mechanisms that drive tumor resistance. By integrating and resolving data from different spatial scales, CORAL can offer novel insights into tumor heterogeneity, and this capability holds tremendous potential for improving our understanding of treatment resistance in cancer, providing critical pathways for future personalized immunotherapies.

## Discussion

CORAL is a deep generative model designed to integrate spatial multioics data using graph aggregation attention mechanisms, enabling comprehensive spatial analysis. The model addresses four key tasks in spatial omics: identifying spatial domains, resolving single-cell components from low-resolution data, predicting cell-cell interactions, and uncovering major molecular regulatory pathways. We demonstrated CORAL’s versatility by benchmarking it against existing methods with overlapping functions. However, no other model achieves all these tasks within a single framework. CORAL’s broad applicability was validated across diverse datasets, including simulated data, datasets directly captured from experimental methods, and synthetic data generated using advanced statistical tools such as scDesign3. Furthermore, we applied CORAL across multiple platforms, including MERFISH, CITE-seq, Stereo-CITE-seq, CODEX, and Visium, showcasing its compatibility and robustness.

The CORAL model showed state-of-the-art advancements in generative modeling by leveraging graph networks and attention layers to conceptualize cells as nodes and their interactions as edges within a graph structure. The model employs contrastive loss and graph Laplacian loss to optimize latent representations, enabling interpretable insights into cellular and molecular relationships in spatial datasets. We believe that these features make CORAL uniquely equipped to address the complexities of spatial omics integration and analysis.

By applying CORAL, we uncovered novel biological insights across multiple datasets. For example, in spatial domain detection, CORAL successfully identified the laminar structure of the mouse cortex, fine-grained structures in the thymus, and heterogeneous tumor microenvironments in hepatocellular carcinoma (HCC). Using CORAL, we uncovered immune-evasive features in non-responders (e.g., P73), including CD8+ T cell exclusion, macrophage recruitment, and enrichment of pathways like JAK-STAT and TGF-*β*. In responders (e.g., P84), reduced immune suppression post-treatment highlighted a more favorable microenvironment. These results demonstrate CORAL’s power to reveal mechanisms underlying therapy outcomes in HCC. In single-cell transcriptomics, CORAL outperformed existing tools, such as SpatialScope, particularly in resolving fine-grained cell types. This capability is especially valuable for spatial proteomics datasets, which often profile a limited number of proteins and lack sufficient resolution for detailed gene ontology analysis. Through high-resolution imputation, CORAL captured novel cell subtypes and enabled downstream analyses equivalent to genome-wide single-cell studies. Additionally, the model elucidated cell-cell interactions, revealing intercellular communication and regulatory networks at unprecedented resolution. In the integration of Visium and CODEX datasets, CORAL harmonized disparate modalities to uncover key biological patterns, further emphasizing its utility for cross-platform analysis.

Despite its strengths, CORAL has certain limitations. The model requires aligned spatial omics data for optimal performance, though perfect alignment or uniform slide dimensions are not strictly necessary. While slight misalignments are permissible, more accurate alignment enhances the reliability and resolution of the results. Addressing these limitations through preprocessing improvements or alignment-free modeling approaches may further expand CORAL’s utility.

Overall, CORAL represents a versatile and powerful tool for spatial omics analysis. Its ability to integrate and resolve data across spatial scales provides unique insights into tissue organization, cellular communication, and molecular regulation. By uncovering novel biological mechanisms and addressing limitations in existing methods, CORAL advances spatial biology and holds tremendous potential for guiding future discoveries and applications in personalized medicine.

## Methods

### The model design

The CORAL model is a deep generative framework designed to integrate multimodal spatial data with varying resolutions. Instead of being limited to specific modalities, CORAL harmonizes data from diverse sources, such as low-resolution spot-level data and high-resolution single-cell data, while preserving spatial structure and modality-specific characteristics. Coral accepts inputs from multiple modalities, denoted as *y*_*ip*_ for single-cell data and *X*_*jg*_ for spot-level data, where *i* ∈ [1, …, *N*] indexes cells, *p* ∈ [1, …, *P*] index single-cell features, *j* ∈ [1, .., *M*] indexes spots, and *g* ∈ [1, …, *G*] indexes spot-level features. Each modality is modeled with a tailored generative process to capture its specific statistical properties.

#### Generative model

CORAL assumes that cells located at spatial coordinates **r**_**i**_ in aligned and adjacent tissue samples share a common underlying latent feature representation, **z**_*i*_ ∈ *ℝ* ^*d*^, which encodes shared biological and spatial information. This latent representation serves as a shared representation across modalities and acts as the foundation for generating both single-cell and spot-level data. The observed data are generated using modality-specific probabilistic distributions, parameterized by neural networks. These distributions are trained to reconstruct observed data while aligning shared latent features in a biologically meaningful way. Specifically: *p*(*y*_*ip*_|**z**_*i*_; *θ*_*y*_) represents the single-cell modality, modeled with a Gamma distribution to capture positive continuous data; *p*(*x*_*ig*_|**z**_*i*_; *θ*_*x*_) represents the inferred single-cell-level data from spot-level observations, modeled with a Negative Binomial distribution to capture overdispersed count data. The observed single-cell and spot-level data are generated using these modality-specific probabilistic distributions:

##### 1. Spot-level modality (*X*_*jg*_)

Spot-level data is modeled with a Negative Binomial distribution to accommodate overdispersed count data:

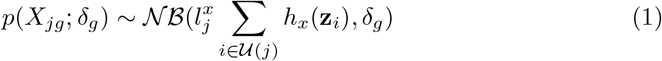

where 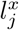 is the library size for spot *j, h*_*x*_ is a neural network parameterizing the rate, 𝒰 (*j*) denotes the set of cells within the spatial neighborhood of spot *j*, and *δ*_*g*_ is the dispersion parameter. Visium spots are larger than single cells, typically covering 1 ∼ 30 cells. Thus, we assume the cells within a circular region |**r**_**i**_ *−* **r**_**j**_| *< ω* are aligned with the spot *j*, and their combined expression generates the observed spot-level data.

##### 2. Single-cell modality (*y*_*ip*_)

Single-cell data follows a Gamma distribution to model positive continuous data:

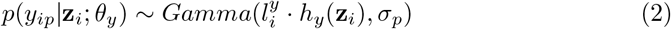

Here, *h*_*y*_ is a neural network parameterizing the shape, 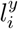 is the library size for cell *i*, and *δ*_*p*_ is the scale parameter.

##### 3. Inferred single-cell representation (*x*_*ig*_)

In addition to the observed modalities, CORAL infers single-cell-level representations *x*_*ig*_ from spot-level data. These representations are also modeled with a Negative Binomial distribution:

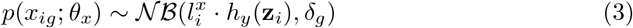

Here, *x*_*ig*_ represents the inferred single-cell RNA-seq expression for cell *i*, reconstructed from spot-level Visium data.

The generative process incorporates a latent variable *z*_*i*_, shared across modalities, and an auxiliary latent variable *v*_*i*_ ∈ *ℝ* ^1^, which disentangles cell state information. The prior distribution of *z*_*i*_ is modeled as:

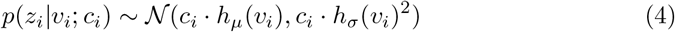

where *c*_*i*_ represents the cell type, *h*_*µ*_ and *h*_*σ*_ are neural networks parameterizing the mean and variance, respectively. The hierarchical latent variable *v*_*i*_ captures additional cell-state-specific variations and is assumed to follow a standard normal prior:

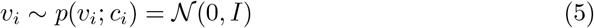

To account for spatial interactions, CORAL constructs a local graph 𝒢 _*i*_ = (𝒱 _*i*_, ℰ_*i*_) for each cell *i*. The graph consists of vertices 𝒱_*i*_, which represent *k* neighboring cells of cell *i*, and ℰ_*i*_ are edges encoding interactions between them. The adjacency matrix *A* ∈0,1^*k×k*^ is defined such that 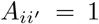 if cell *i* and cell *i*^*′*^ are neighbors, and 0 otherwise. The spatial interactions between cells are captured by the probability of an edge between two nodes:

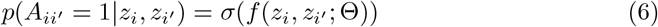

Where *σ* is the sigmoid function and *f* is a neural network parameterized by Θ. The full generative model is expressed as *p*(**X, y**, 𝒜, *x, v, z*|*c*), with equals to,

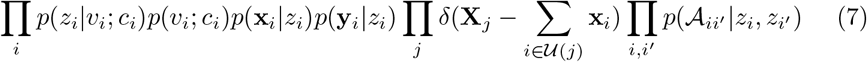

Each component is defined as:

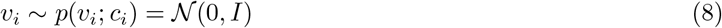

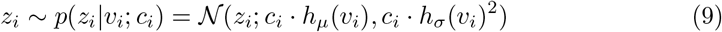

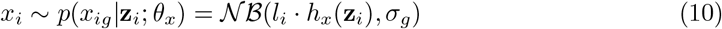

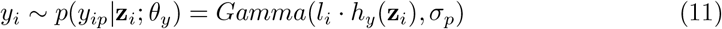

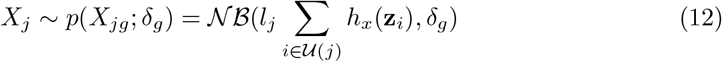

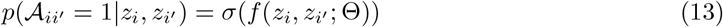

#### Inference model

CORAL employs a joint variational inference model to approximate the posterior distributions. The joint posterior is expressed as:

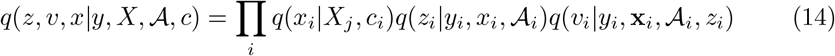

The major items include:

1. **Single-cell inference** *q*(*x*_*i*_|*X*_*j*_, *c*_*i*_) Infers the single-cell expression profile **x**_*i*_ conditioned on the spot-level modality data *X*_*j*_ and the cell type *c*_*i*_.
2. **Latent spatial feature inference** *q*(*z*_*i*_|*yi, xi*, 𝒜_*i*_) Approximates the posterior distribution of the latent feature vector *z*_*i*_ for cell *i*, considering the cell’s observed single-cell modality data **y**_*i*_, its inferred RNA-seq expression **x**_*i*_, and the local graph 𝒢_*i*_ representing spatial and potentially functional relationships with neighboring cells. This implies that the latent features *z*_*i*_ are influenced by both the molecular data and the cell’s spatial context within its microenvironment.
3. **Latent spatial feature inference** *q*(*v*_*i*_|**y***i*, **x***i*, 𝒢_*i*_, *z*_*i*_) Captures the major compnents of spatial specific states, integrating single-cell modality data **y**_*i*_, its inferred expression **x**_*i*_, the local spatial graph 𝒢_*i*_, and the latent features *z*_*i*_. Each posterior distribution is parameterized using neural networks, ensuring flexible and efficient inference.
4. Graph attention networks for modeling cell-cell interations To model the cell-cell interactions, CORAL incorporates Graph Attention Networks (GATs). GATs use an attention mechanism to dynamically weigh neighbor contributions. The attention coefficient is defined as:

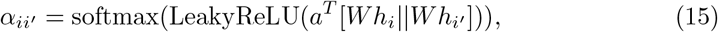

where is a learnable weight vector, *W* is a weight matrix applied to the node features *h*, and || denotes concatenation. The attention coefficients influence how neighboring cells’ features are aggregated, enabling CORAL to capture spatially dependent interactions.

#### Learning and optimization

CORAL optimizes its parameters by maximizing the evidence lower bound (ELBO),which integrates reconstruction accuracy and prior regularization, while capturing the hierarchical relationships between spot-level data (*X*) and singl-cell-level data (*x*). The ELBO is defined as:

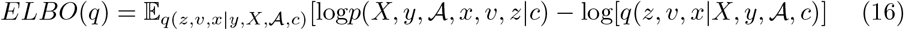

This formulation reflects the trade-off between accurately modeling the observed data and ensuring the learned latent variables align with the prior distributions. The joint probability *p*(*X, y*, 𝒜, *x, v, z*|*c*) represents the generative process of the model, and the posterior *q*(*z, v, x*|*X, y*, 𝒜, *c*) is approximated using neural networks. The loss function incorporates reconstruction terms, regularization, and spatial constraints to ensure accurate modeling of multimodal data, alignment of latent variables with biological context, and preservation of spatial relationships.

1. **Reconstruction Terms:**

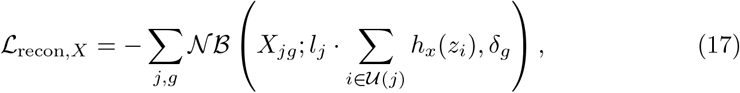

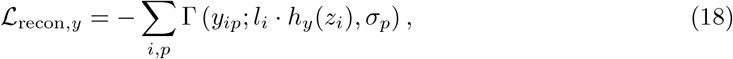

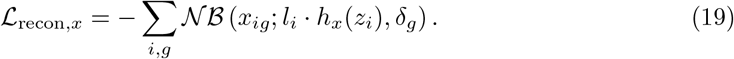 These reconstruction terms ensure that the observed data (*X, y*) and inferred singlecell data (*x*) are faithfully reproduced by the generative model.
2. **KL Divergence**

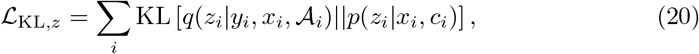

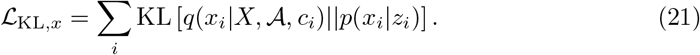 The KL divergence terms regularize the latent variables, aligning the approximate posterior distributions with their respective priors.
3. **Graph Laplacian Regularization:**

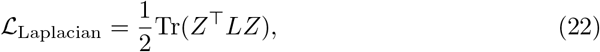

where *Z* represents latent embeddings, and *L* is the graph Laplacian matrix. This term enforces smoothness among latent variables for spatially adjacent cells, encouraging biologically meaningful relationships between neighboring cells.
4. **Contrastive Loss:**

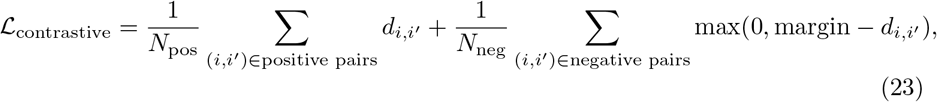

where 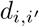 is the distance between latent representations. The contrastive loss enhances the separation between biologically distinct cell types (negative pairs) while preserving similarity among the same types (positive pairs). This ensures that the latent embeddings are biologically interpretable.

The total loss is a weighted combination of all components, where the hyperparameters *λ*_*X*_, *λ*_*y*_, *λ*_*x*_, *λ*_*z*_, *λ*_Lap_, and *λ*_contrast_ balance their contributions. This comprehensive loss function ensures robust integration of multimodal data, accurate reconstruction, and biologically meaningful latent representations.

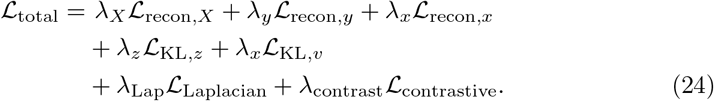

CORAL is trained with the Adam optimizer using gradient clipping to stabilize updates. A dynamic learning rate scheduler adjusts the learning rate, ensuring convergence. The model iteratively updates parameters by reconstructing multimodal data, aligning latent variables, and enforcing spatial constraints. By combining these components, CORAL produces robust, biologically meaningful representations.

### Cluster Separation Ratio

The cluster separation ratio quantifies the quality of clustering in the principal component (PC) space by comparing the compactness of points within clusters (intra-cluster distances) to the separation between clusters (inter-cluster distances). It is defined as:

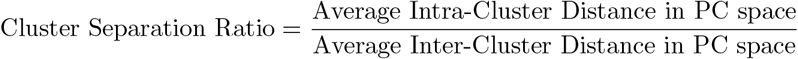

Where: Intra-Cluster Distance is the average pairwise distance between all points within the same cluster:

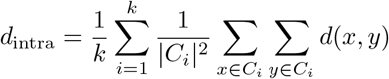

Here, *k* is the number of clusters, *C*_*i*_ represents the set of points in cluster *i*, and *d*(*x, y*) is the distance between points *x* and *y*.

Inter-Cluster Distance is the average pairwise distance between the centroids of different clusters:

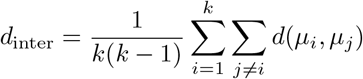

Here, *µ*_*i*_ and *µ*_*j*_ are the centroids of clusters *i* and *j*, respectively.

A lower cluster separation ratio indicates better-defined clusters, as it suggests that points within clusters are tightly packed (low *d*_intra_) and well-separated from points in other clusters (high *d*_inter_).

## Supplementary information

Figures S1-S9

## Acknowledgements

We thank MDACC for providing HCC data.

## Author contributions

J.Zou and A.T. conceived the study and provided overall supervision of the study. S.H. conceived, designed and developed CORAL. S.H., M.B., D.S., J.Zhou analyzed and interpreted data. S.Q. provided additional supervision. S.H., M.B., D.S., J.Zhou, B.C., J.B., Z.W., S.Q. A.T. and J.Zou wrote the paper. All authors reviewed, contributed to and approved the paper.

## Data availability

The synthetic datasets used in this study, provided by SpatialGlue, can be accessed at https://zenodo.org/records/7879713#.ZE3aOnZByUk. The public MERFISH dataset was obtained from https://github.com/YangLabHKUST/SpatialScope. Synthetic data generated from CITE-seq data using scDesign3 is available via https://figshare.com/ndownloader/files/40581968. The Stereo-CITE-seq mouse thymus data, also provided by SpatialGlue, is accessible at https://zenodo.org/records/7879713#.ZE3aOnZByUk. The raw single-cell sequencing data of HCC generated in this manuscript will be deposited in the National Center for Biotechnology Information’s Gene Expression Omnibus.

## Code availability

The CORAL package and code to reproduce the results in this manuscript is available on the GitHub repositories (https://github.com/zou-group/CORAL).

**Supplementary Fig. 1:**
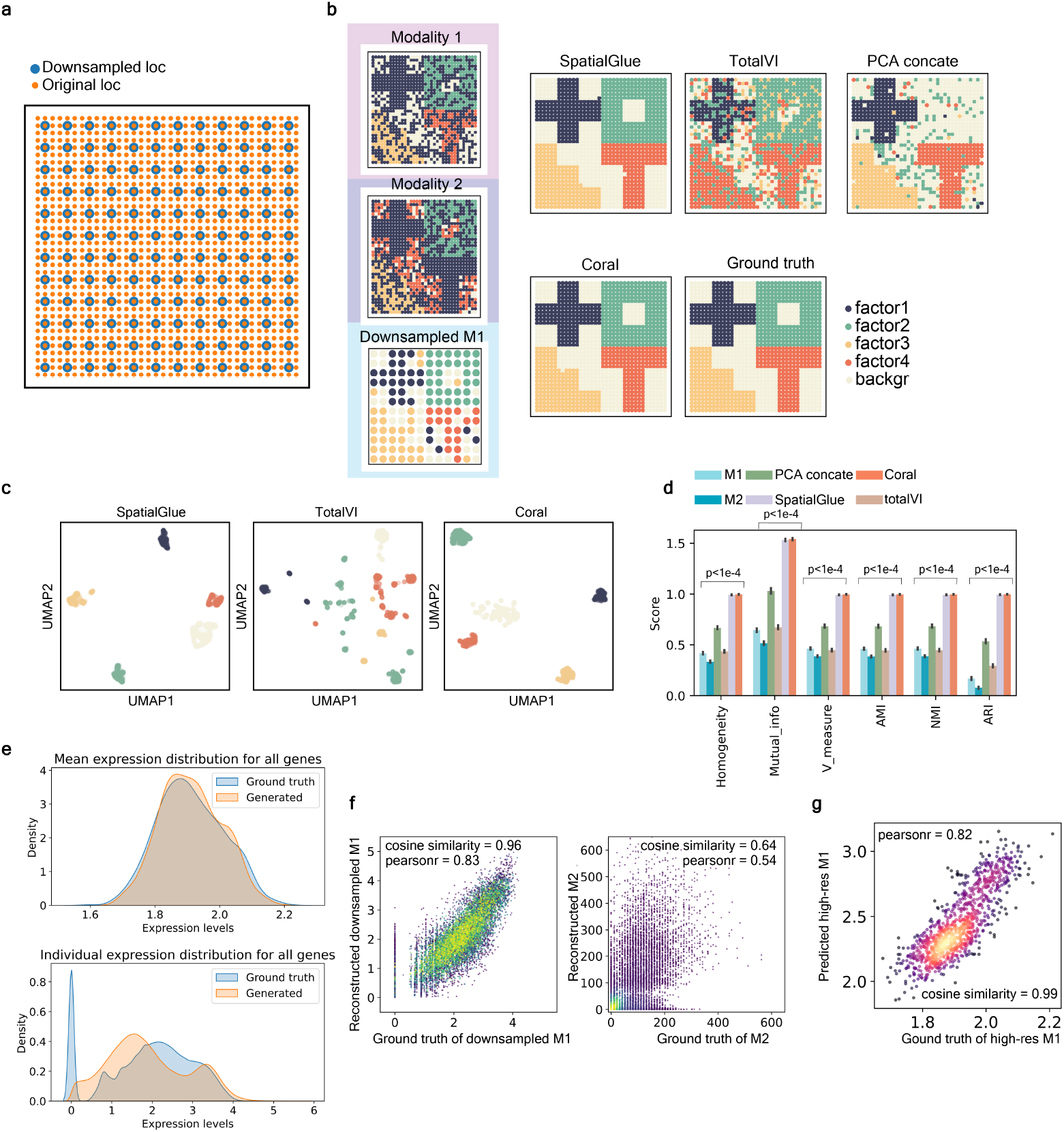
CORAL applied on synthetic data. The data simulated processes was described in the SpatialGlue[22]. **a**, Location of downsampled and original spots. **b**, Left: leiden clustering of modality 1, modality 2 and downsampled modality 1. Right: leiden clustering on latent variables identified by SpatialGlue, TotalVI, CORAL, concatenating PCA of two modality, compared with ground truth. **c**, UMAP OF latent variables identified by SpatialGlue, TotalVI, and CORAL. **d**, Performance metrics (Homogeneity, Mutual Information, V-measure, Adjusted Mutual Information (AMI), Normalized Mutual Information (NMI), Adjusted Rand Index (ARI)) comparing CORAL’s spatial domain detection accuracy with modality 1, modality2, Spatial Glue, totalVI, and concatenated PCA values of two modalities. The one-way Anova test was performed on each metric, p value *<* 1e-4 for all tests. **e**, Upper: distribution of mean expression between all genes generated by CORAL and ground truth. Lower: distribution of individual expression between all genes generated by CORAL and ground truth. **f**, Scatter plot of reconstructed and ground truth of expression of individual genes in downsampled modality 1 and modality 2, with cosine 23 similarity 0.96, and 0.64 separately, and pearson correlation 0.83 and 0.54. **g**, Scatter plot of predicted mean expression of individual genes for high-resolution modality 1 and ground truth modality 1. Pearson correlation is 0.82, and cosine similarity is 0.99.

**Supplementary Fig. 2:**
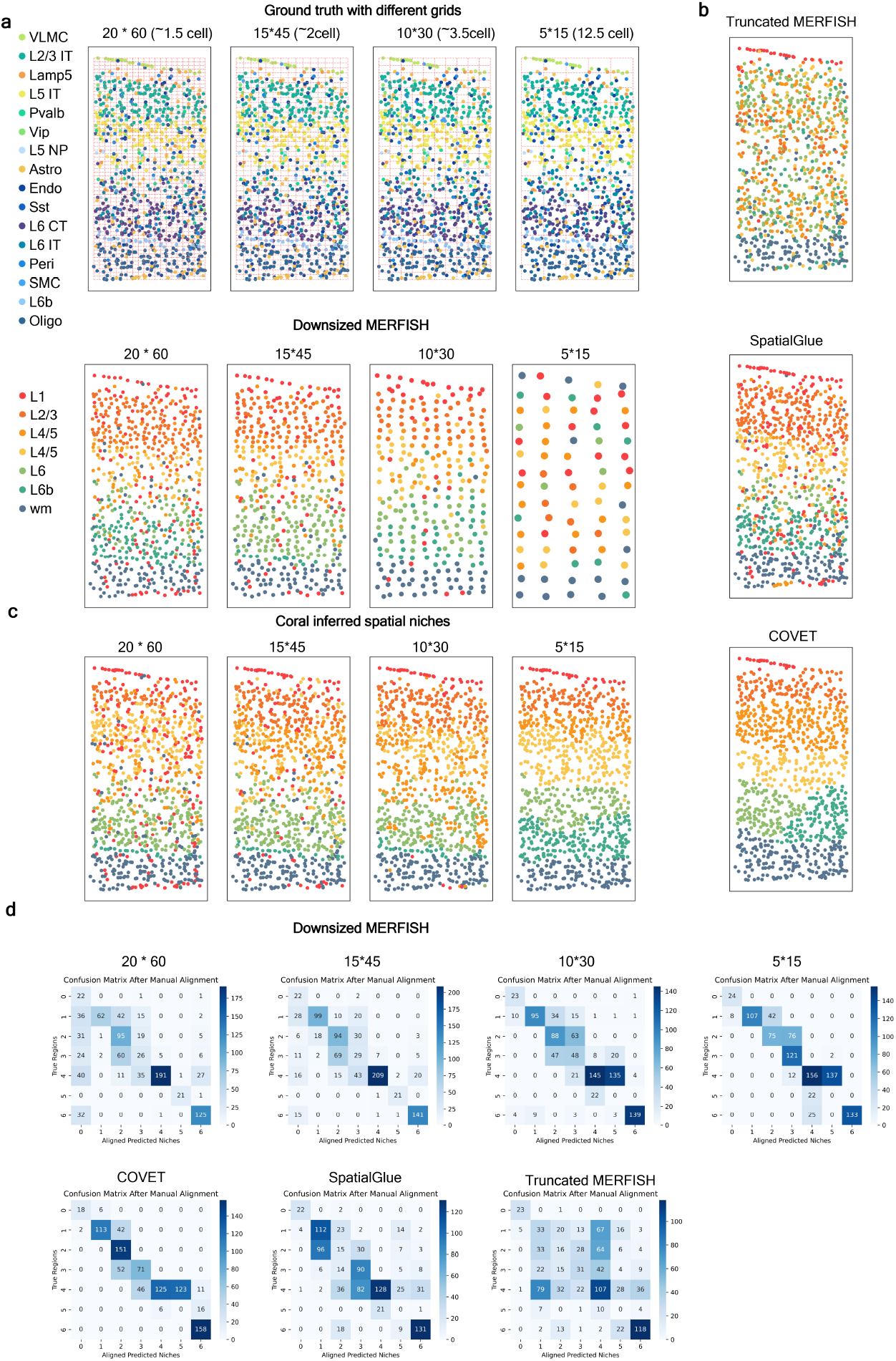
CORAL applied to simulated MERFISH data. **a**, Ground truth spatial niches with different grid resolutions (20×60, 15×45, 10×30, and 5×15) simulating varying cell densities. Each point represents a cell, color-coded by its true niche. **b**, Truncated MERFISH and SpatialGlue inferred niches, illustrating the impact of downsampling and alignment quality on niche identification. **c**, Spatial niches inferred by CORAL at varying grid resolutions (20×60, 15×45, 10×30, 5×15). CORAL demonstrates robustness to spatial resolution changes while maintaining biological consistency in niche assignments. **d**, Confusion matrices for the predicted spatial niches compared to ground truth after manual alignment. Top row: Downsampled MERFISH data at different resolutions. Bottom row: Comparison of CORAL with COVET, SpatialGlue, and Truncated MERFISH metho2d4s. CORAL achieves high accuracy in niche alignment, as indicated by the diagonal dominance in confusion matrices.

**Supplementary Fig. 3:**
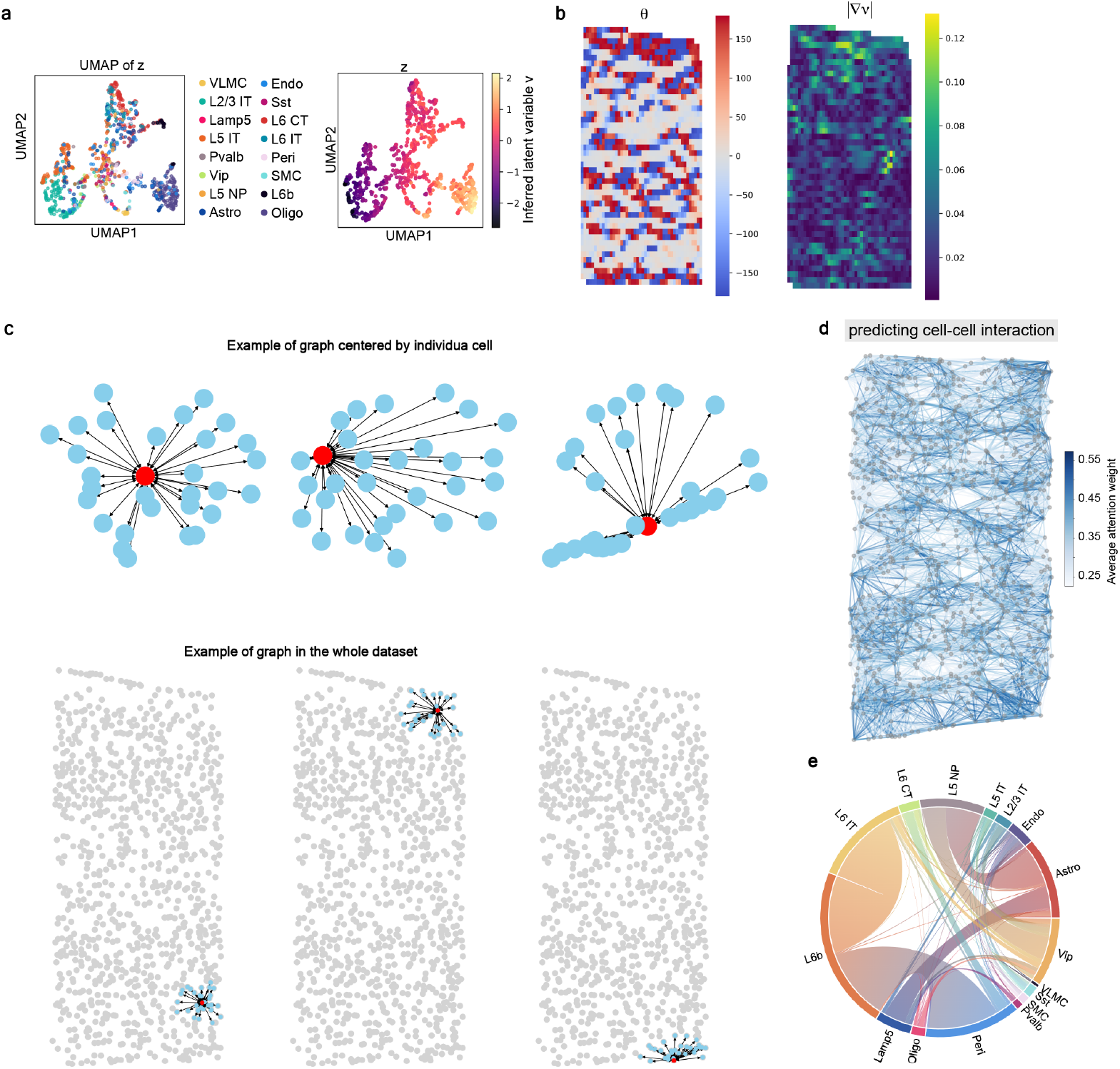
Graphical and latent representation analysis in CORAL. **a**, UMAP visualizations of truncated MERFISH data, latent representation *z*, and latent representation *v*. The latent space aligns well with biological cell types (color-coded). **b**, Heatmaps of *θ* (left) and |∇*v*| (right), highlighting latent spatial patterns and gradients of the hierarchical latent variable *v*. **c**, Graph-based visualization of spatial relationships captured by CORAL. Top row: examples of graphs centered on individual cells, showing local neighborhoods. Bottom row: examples of spatial graphs in the entire dataset. CORAL effectively encodes spatial connectivity, preserving local and global tissue structures. **d**, Network map of cell-cell interactions across the mouse motor cortex, with color intensity representing interaction strength. **e**, Circular plot of cell-type interactions, where bar lengths represent total mean interactions and band widths represent interaction strength.

**Supplementary Fig. 4:**
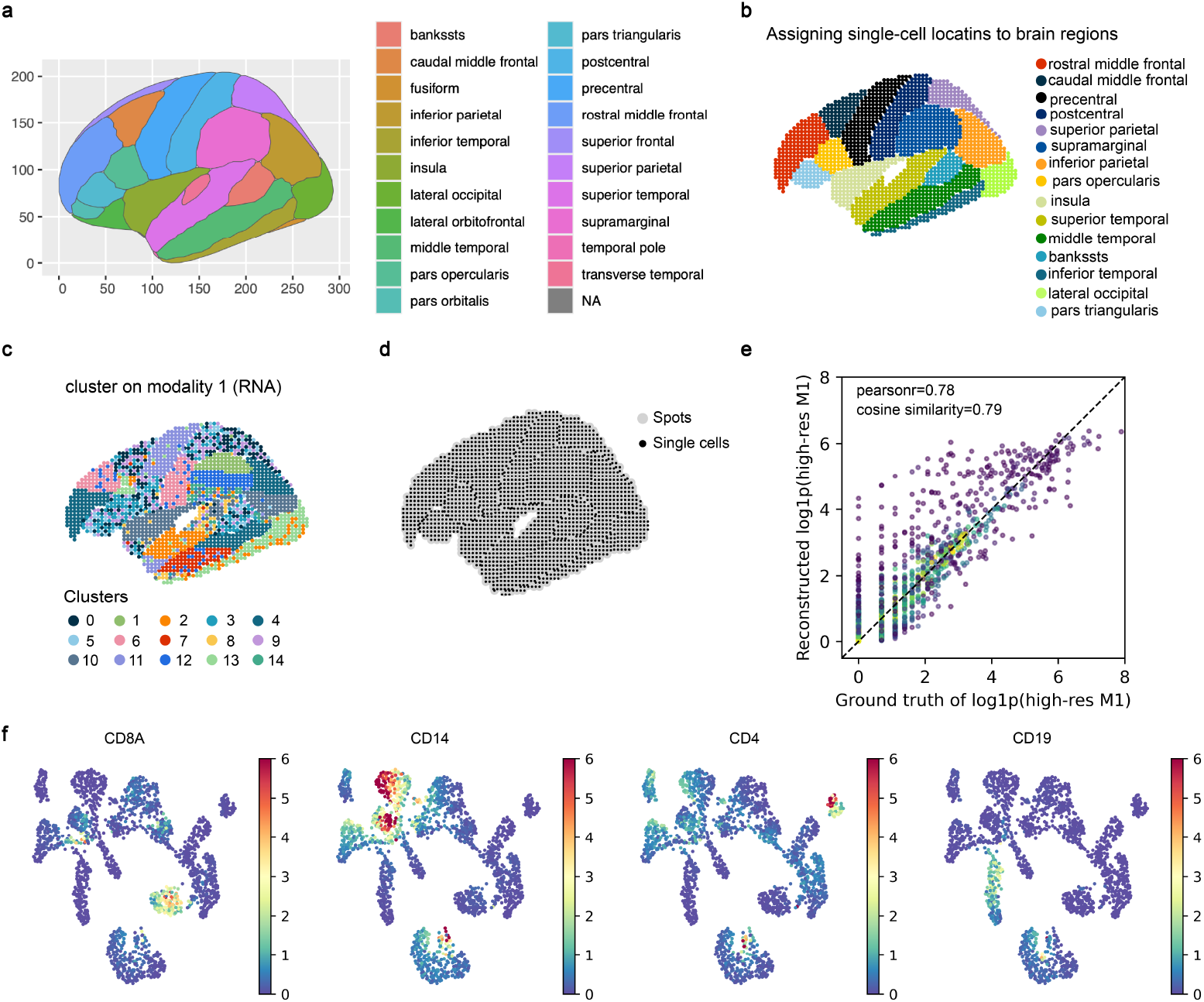
CORAL applied to sythetic brain spatial transcriptomics data. **a**, Annotated brain regions, with each color representing a distinct anatomical region. **b**, Assignment of single-cell locations to brain regions based on CORAL’s spatial alignment, using both high-resolution and low-resolution modalities. **c**, Clustering of single cells based on RNA modality (modality 1), revealing spatially coherent biological clusters across the brain regions. **d**, Visualization of single-cell and spot-level data spatial locations, showing the integration of multiple resolutions. **e**, Scatter plot comparing reconstructed high-resolution modality 1 (*M*_1_) with ground truth expression values. CORAL achieves a Pearson correlation of 0.78 and a cosine similarity of 0.79, indicating high fidelity in reconstructing single-cell expression data. **f**, Heatmaps showing the expression of marker genes (CD8A, CD14, CD4, CD19) across the integrated dataset, highlighting CORAL’s ability to preserve spatial and molecular patterns at single-cell resolution.

**Supplementary Fig. 5:**
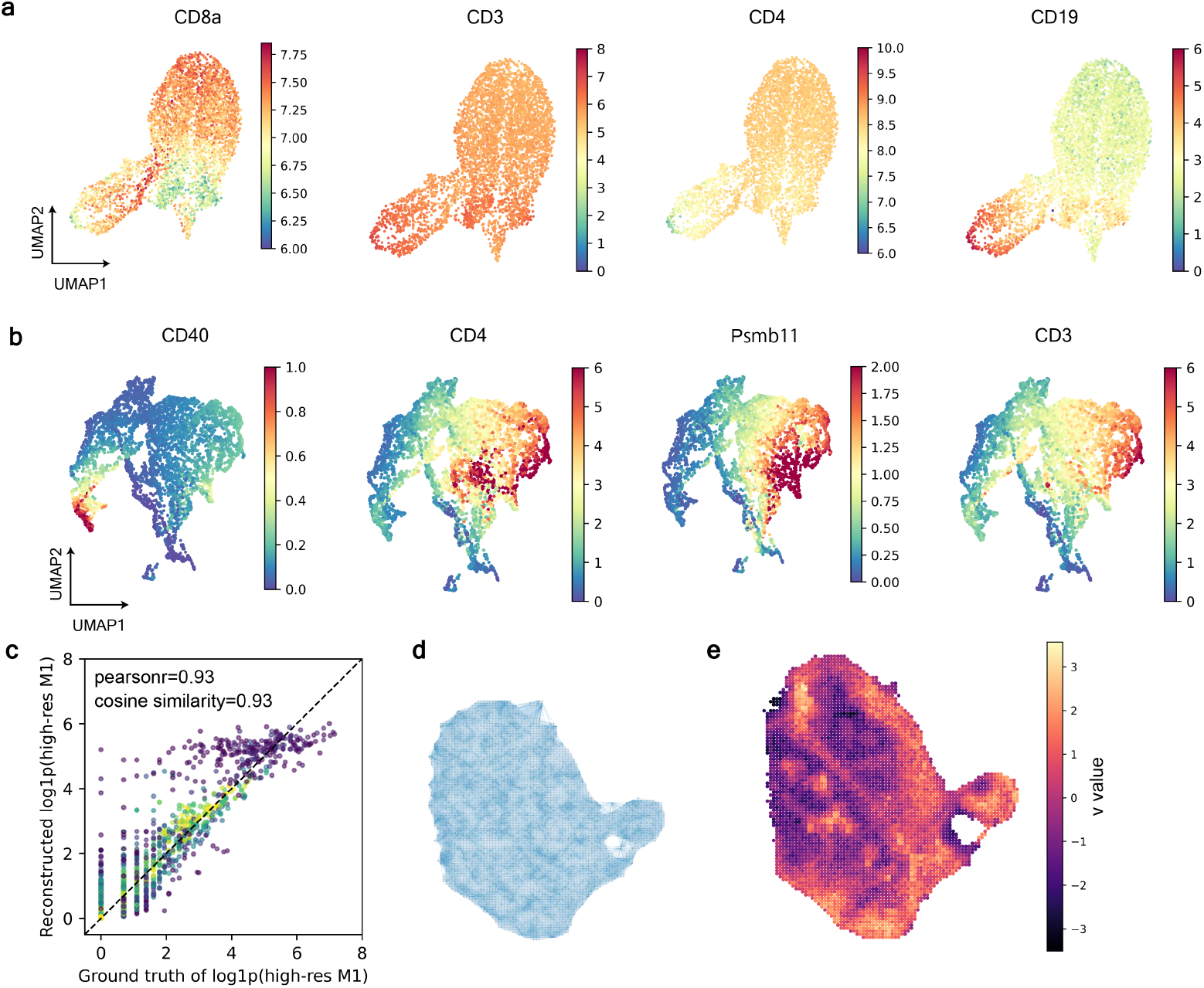
CORAL preserves spatial and molecular patterns in immune datasets. **a**, UMAP visualizations of marker gene expressions (CD8A, CD3, CD4, CD19) in the integrated dataset. CORAL successfully captures distinct immune cell subtypes in the latent space. **b**, Heatmaps of additional marker genes (CD40, CD4, Psmb11, CD3) in the UMAP representation, highlighting CORAL’s ability to preserve gene-specific patterns. **c**, Scatter plot comparing reconstructed high-resolution modality 1 (*M*_1_) with ground truth expression values. CORAL achieves a Pearson correlation of 0.93 and cosine similarity of 0.93, demonstrating high accuracy in singlecell expression reconstruction. **d**, Visualization of the spatial locations of cells in the dataset, showing the reconstructed tissue structure. **e**, Heatmap of the *v*-value latent variable over the spatial domain, highlighting spatial gradients captured by CORAL.

**Supplementary Fig. 6:**
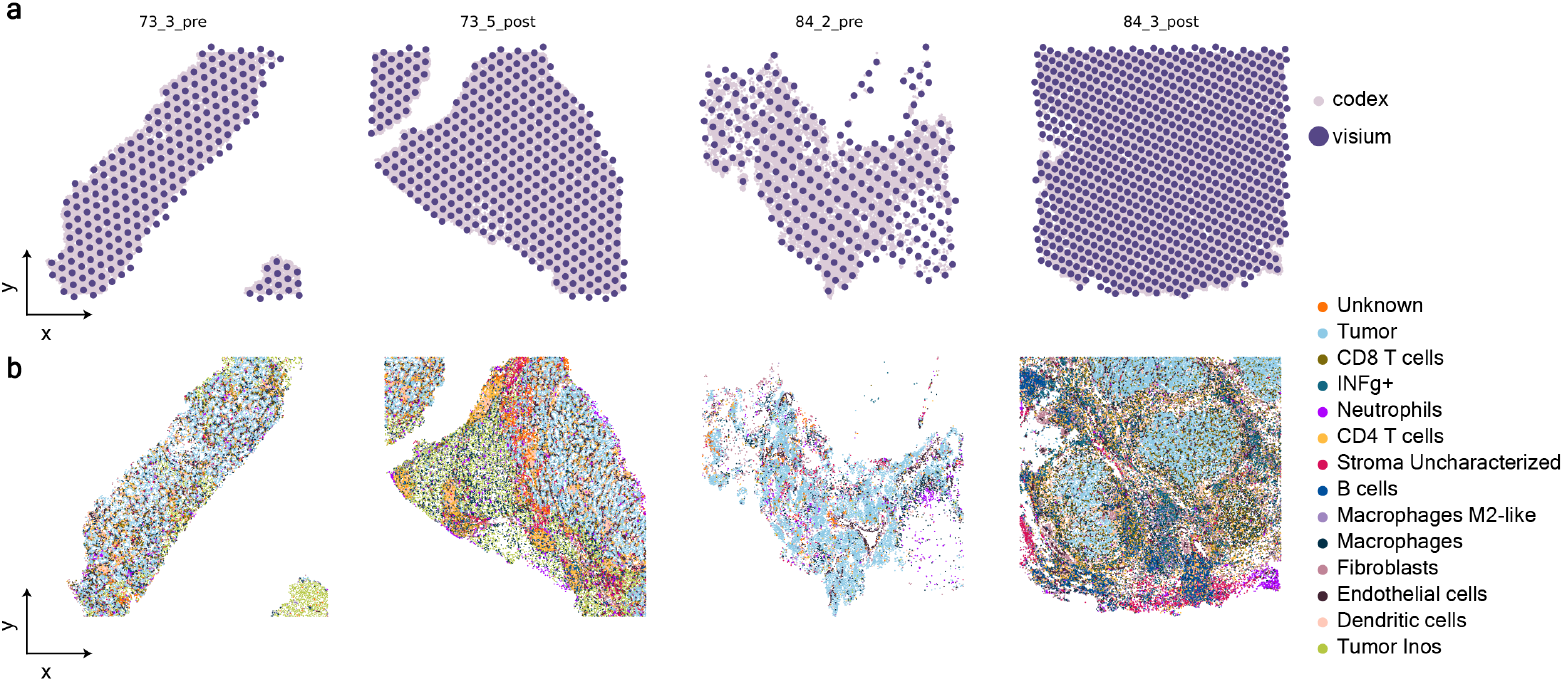
Spatial integration of CODEX and Visium data using CORAL. **a**, Spatial distribution of Visium and CODEX datasets for four samples (73_3_pre, 73_5_post, 84_2_pre, and 84_3_post). Visium and CODEX spots are overlaid to demonstrate the integration of multi-modal spatial data. **b**, Cell type annotation across the integrated dataset, with color-coded cell types including tumor cells, immune cells (CD8 T cells, CD4 T cells, B cells, neutrophils, macrophages), stromal cells (fibroblasts, endothelial cells), and dendritic cells. CORAL successfully aligns spatial and cell type data, capturing the complex tumor microenvironment.

**Supplementary Fig. 7:**
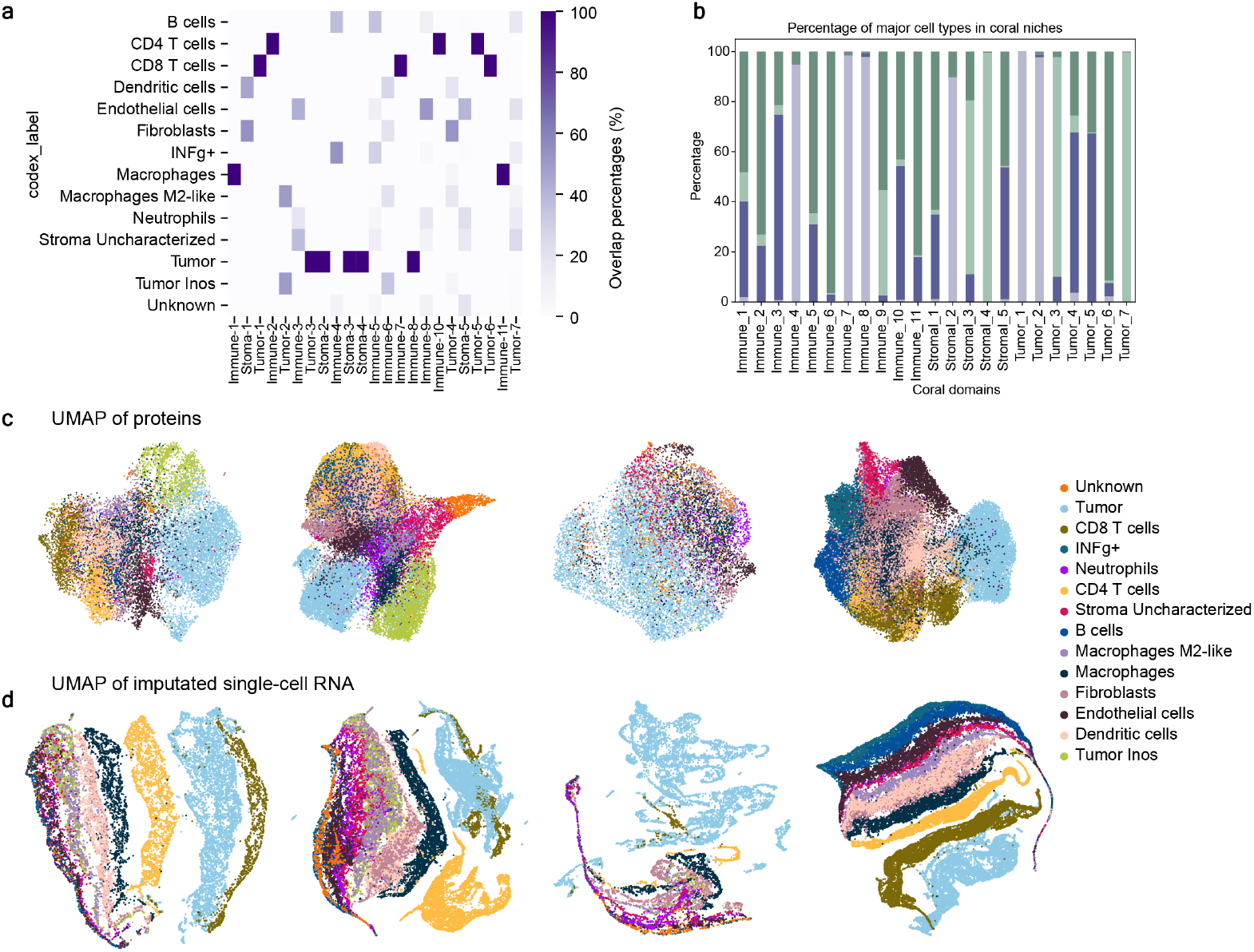
Cell type composition and latent space visualization in CORAL niches. **a**, Heatmap showing the overlap percentage between cell types in the CODEX dataset and CORAL-inferred spatial domains. The strong diagonal highlights accurate cell type assignments within CORAL niches. **b**, Bar plot showing the percentage composition of major cell types (e.g., immune, stromal, tumor) across CORAL-inferred domains. The results demonstrate that CORAL effectively segregates niches based on their dominant cell type composition. **c**, UMAP visualization of protein expression levels in the integrated dataset, with distinct clusters corresponding to cell types. **d**, UMAP visualization of imputed single-cell RNA expression from CORAL, showing spatial and molecular alignment with cell types identified in the CODEX dataset. CORAL preserves the spatial heterogeneity of the tumor microenvironment while capturing fine-grained molecular patterns.

**Supplementary Fig. 8:**
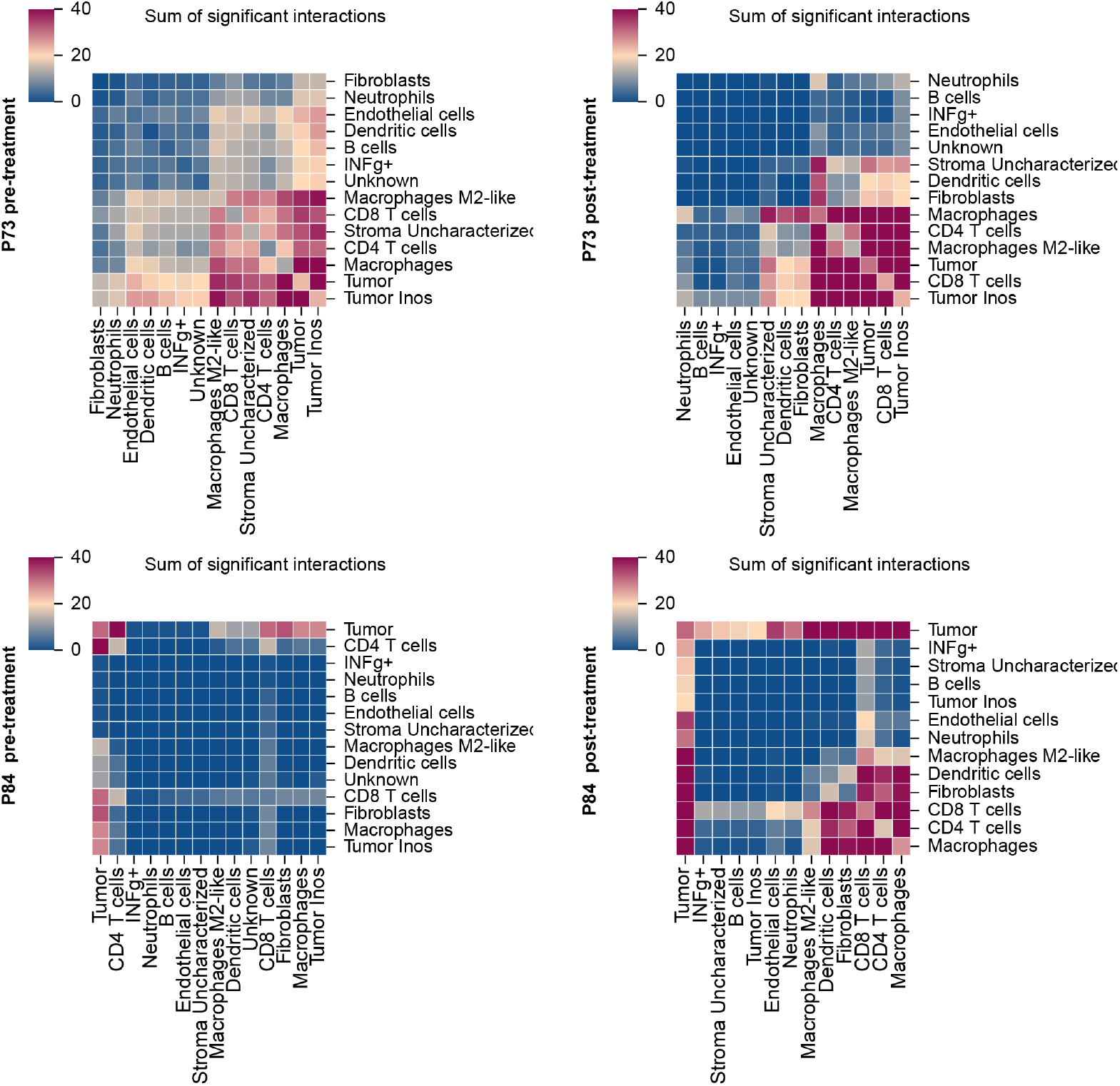
Cell-cell interaction analysis pre- and post-treatment using CORAL. Heatmaps showing the sum of significant interactions between cell types across different treatment conditions for samples P73 and P84. Each cell type is represented along the rows and columns, and the intensity of the heatmap indicates the interaction strength. **a**, Sample P73 pre-treatment, highlighting strong interactions between tumor cells, macrophages, and CD4 T cells. **b**, Sample P73 post-treatment, showing changes in interaction patterns, including increased interactions involving tumor-infiltrating immune cells. **c**, Sample P84 pre-treatment, with significant interactions dominated by tumor cells, stromal cells, and CD8 T cells. **d**, Sample P84 post-treatment, indicating shifts in cell-cell interactions, particularly involving macrophages and stromal cell populations. CORAL effectively captures dynamic changes in the tumor microenvironment associated with treatment response.

**Supplementary Fig. 9:**
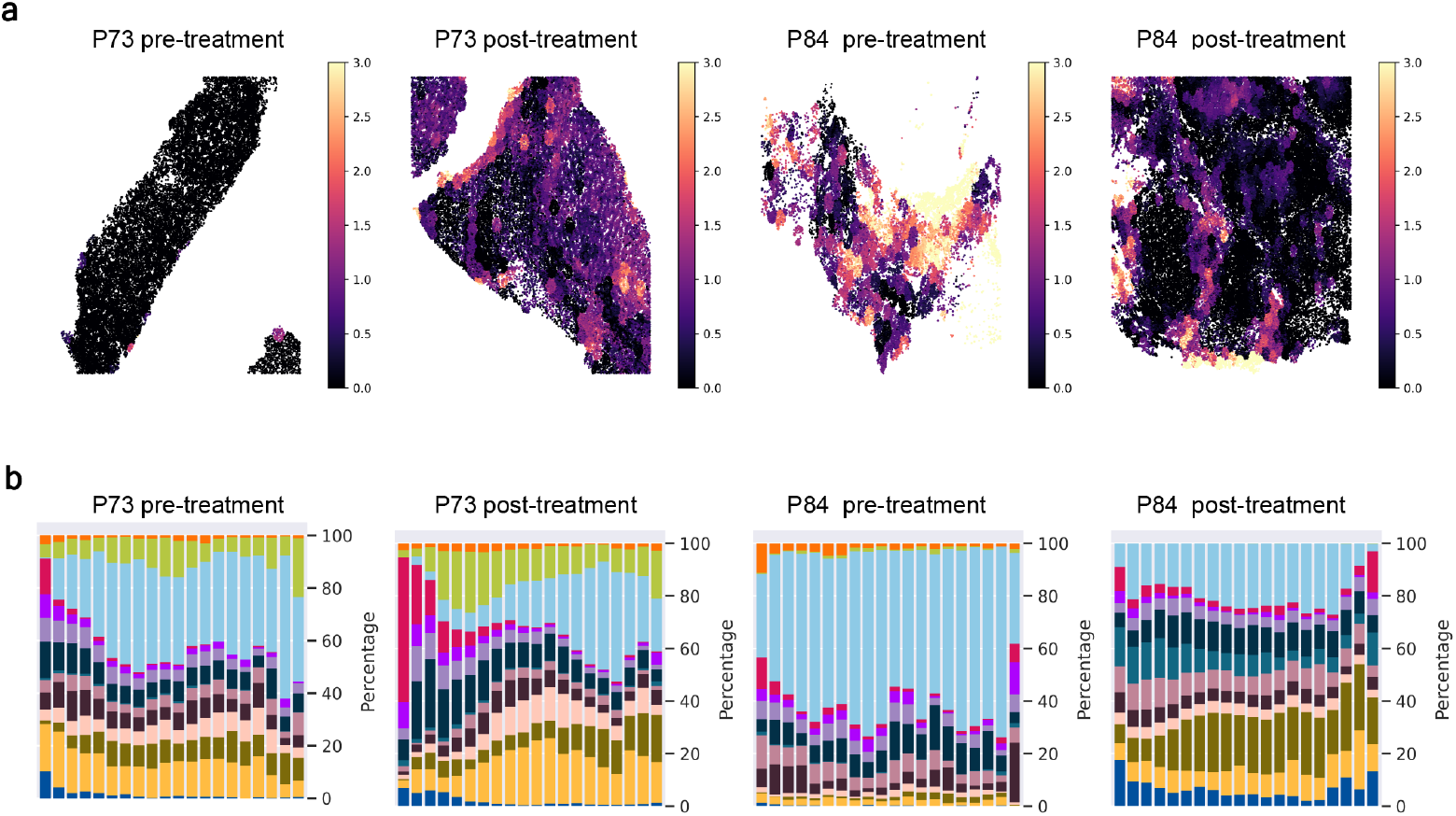
Spatial patterns and cell type composition pre- and post-treatment. **a**, Heatmaps of the spatial *v*-value latent variable for samples P73 and P84 under pre-treatment and post-treatment conditions. The *v*-values reveal spatial gradients and shifts in tissue microenvironments, capturing treatment-induced changes in tumor and immune interactions. **b**, Stacked bar plots showing the percentage composition of major cell types across spatial niches for each condition. Each bar represents a spatial niche, and colors correspond to cell types, including tumor cells, immune cells (e.g., CD4 T cells, CD8 T cells, B cells), and stromal cells. The plots highlight changes in cell type proportions associated with treatment responses, such as shifts in immune cell infiltration and tumor composition.

